# Ribosomal Protein SA (RPSA) is required for localized translation and sarcomere maintenance in mice

**DOI:** 10.1101/2023.07.26.550187

**Authors:** Rami Haddad, Omer Sadeh, Tamar Ziv, Itai Erlich, Lilac Haimovich-Caspi, Ariel Shemesh, Jolanda van der Velden, Izhak Kehat

**Author notes:** Authorship note: RH and OS contributed equally to this work. Correspondence: Izhak Kehat MD,PhD The Ruth and Bruce Rappaport Faculty of Medicine Technion-Israel Institute of Technology Efron St. no number P.O.B. 9649 Bat Galim, Haifa 31096, Israel.

## Abstract

Cardiomyocyte sarcomeres contain localized ribosomes, but the factors responsible for their localization and the significance of localized translation are unknown. Using proximity labeling, we identified Ribosomal Protein SA (RPSA) as a Z-line protein. In cultured cardiomyocytes, the loss of RPSA led to impaired local protein translation and reduced sarcomere integrity. By employing CAS9 expressing mice along with adeno-associated viruses expressing CRE recombinase and single-guide RNAs targeting *Rpsa*, we knocked out *Rpsa* in vivo and observed mis-localization of ribosomes and diminished local translation. These genetic mosaic mice with *Rpsa* knockout in a subset of cardiomyocytes developed dilated cardiomyopathy, featuring atrophy of RPSA-deficient cardiomyocytes, compensatory hypertrophy of unaffected cardiomyocytes, left ventricular dilation, and impaired contractile function. We demonstrate that RPSA C-terminal domain is sufficient for localization to the Z-lines and that if the microtubule network is disrupted RPSA loses its sarcomeric localization. These findings highlight RPSA as a ribosomal factor essential for ribosome localization to the Z-line, facilitating local translation and sarcomere maintenance.

## Introduction

Local translation has several benefits over diffuse or unlocalized protein production. Importantly, it allows protein synthesis closer to the site of action, is more economic, may prevents protein aggregation, and can facilitate processes such as co-translational assembly (1). Local translation requires the presence of the mRNA, the ribosome, as well as additional translation factors and co-factors. While the localization of mRNAs and the RNA-binding protein involved, particularly in neurons, are well studied (2), the factors involved in the localization of cytoplasmic ribosomes are less clear.

The sarcomere is the basic contractile element of cardiomyocytes. Because its protein constituents have a limited half-life, this complex must be continuously maintained (3). Failure in sarcomeric proteostasis may lead to heart failure because failing human hearts have reduced sarcomeres and sarcomeric protein content (4–6). We previously showed that ribosomes are localized on both sides of the Z-line of the sarcomere, and that proteins are translated there, leading us to propose that sarcomeres are maintained by the localized translation of proteins within them (7). We have also demonstrated that the microtubule cytoskeleton and kinesin are essential for localized translation in the heart (7, 8). Similar observations were made for skeletal muscles (9). Yet, the importance of localized translation for sarcomere maintenance has not been demonstrated, and the ribosomal factors required for localization are unknown.

Here we used proximity labeling to elucidate the proteomic composition of sarcomeric Z-lines in an unbiased manner and identified Ribosomal Protein SA (RPSA), a protein with reported ribosome and laminin receptor functions. We show that loss of RPSA in cultured cardiomyocytes results in the loss of local protein translation and in reduced sarcomere integrity. Using conditional and unconditional CRISPR associated protein 9 (CAS9) expressing mice and adeno-associated viruses (AAVs) expressing CRE recombinase in cardiomyocytes with single-guide RNAs (sgRNAs) targeting *Rpsa* we knocked out *Rpsa* in cardiomyocytes in vivo and show that the loss of RPSA results in mis-localization of ribosomes and reduced local protein translation. These genetic mosaic mice, where only a portion of cardiomyocytes are transduced and have *Rpsa* knockout, develop a dilated cardiomyopathy with atrophy of *Rpsa* knockout cardiomyocytes, compensatory hypertrophy of untransduced cardiomyocytes, left ventricular dilation and reduced contractile function. RPSA is known to directly bind the microtubular system and we show its C-terminal domain is sufficient for localization to Z-lines, and that if the microtubule network is disrupted RPSA loses its sarcomeric localization. These data show RPSA, the first ribosomal factor responsible for localizing cytoplasmic ribosomes identified, is required for localization of ribosomes to the Z-line, for the local translation there, and for the maintenance of the sarcomere.

## Results

### Proximity-dependent labeling identifies components of translational machinery at sarcomeric Z-lines

To investigate the proteomic composition of sarcomeric Z-lines and identify the factors involved in localized translation, we employed the BioID2 proximity labeling assay using Cypher (LDB3), a known Z-line protein, as bait. For this purpose, we generated a fusion construct between Cypher and the promiscuous biotin ligase BioID2. We then transduced neonatal rat ventricular cardiomyocytes (NRVMs) with an adenoviral vector encoding for Cypher-BioID2 or control virus expressing BioID2 alone. To confirm the correct subcellular localization of our fusion construct we first performed immunofluorescence staining that demonstrated that Cypher-BioID2 fusion construct closely matched the known localization pattern of endogenous Cypher at the sarcomeric Z-lines, whereas BioID2-alone displayed a more diffuse cytoplasmic staining (Figure 1A). We then used three independent samples for each group, each containing 12 million cardiomyocytes, and purified biotinylated proteins using streptavidin-coated magnetic beads after 18 hours of incubation with biotin. Western blot analysis of the extracted cardiac proteins, using a streptavidin probe confirmed the successful biotinylating of proteins and showed distinct sets of labeled proteins in the Cypher-BioID2 sample compared to the BioID2-alone control samples (Figure 1B).

**Figure 1.**
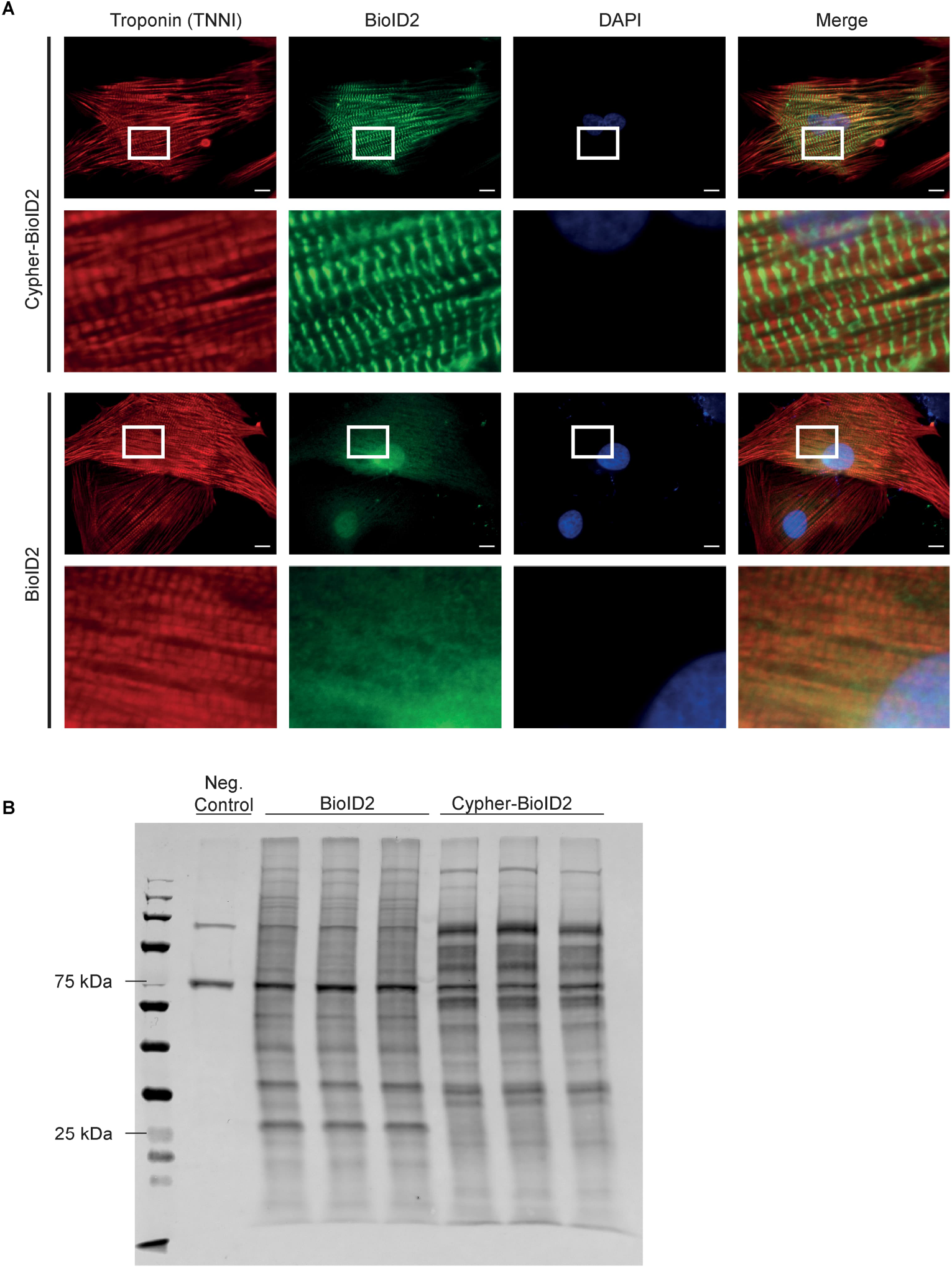
Cypher-BioID2 is localized to the Z-lines and biotinylates specific proteins. (**A**) Representative immunofluorescence images of NRVMs transduced with adenoviruses encoding for either Cypher-BioID2 (bait construct) or BioID2 (control construct) with inset showing Z-line localization of Cypher-BioID2 and a diffuse cytoplasmic localization for BioID2 alone. Samples were immunostained for cardiac troponin (red), BioID (green) and DAPI (blue). Scale bar = 10 µm. (**B**) Western blot analysis of biotinylated proteins using IRDye800-Streptavidin showing different biotinylated patterns for cardiomyocytes transduced with BioID2 or Cypher-BioID2 and minimal biotinylation of the untransduced negative control sample. Each lane was loaded with 100 µg of protein lysate.

Pulled down proteins were trypsin digested on beads and subjected to mass spectrometry analysis. The proteomic analysis identified 135 proteins significantly enriched in the Cypher-BioID2 sample compared to the BioID2-alone control using a cutoff of more than 2-fold enrichment (log2 FC≥1) and t test p value <0.05 (Figure 2A, Table 1). As expected, LDB3 (Cypher), that served as the bait, was the most abundant and most significantly enriched protein in the bait group. Proteins known to be localized to the Z-line, such as α-actinin (ACTN2) and titin (TTN), were also significantly enriched in the bait group. On the other hand, highly expressed proteins known to be localized to the M-line such as MYOM1 (Myomesin1) or OBSCN (Obscurin) were significantly more abundant in the control group, demonstrating we have specifically enriched for a Z-line nanodomain. Additional proteins significantly enriched in the bait group included RPSA (Ribosomal Protein SA), and RPL38 (Ribosomal Protein L38).

**Figure 2.**
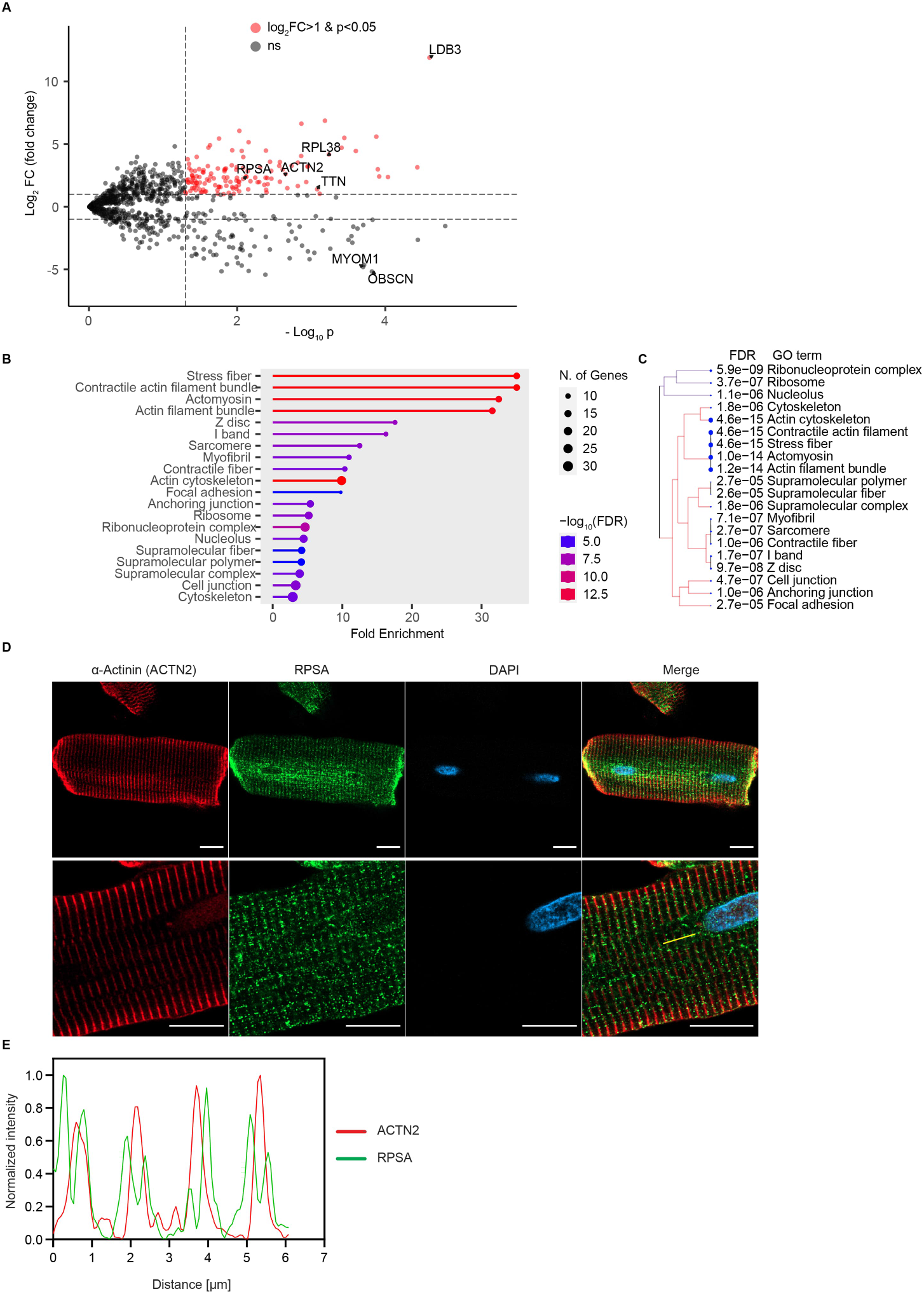
Proximity-labeling with Cypher-BioID2 identifies Z-line proteins and components of the translational machinery. (**A**) Volcano plot of proteins identified by mass spectrometry showing the log2 fold change in protein abundance with Cypher-BioID2 vs. BioID2 alone and the t-test -log10 (p value) of the difference significance. Dashed horizontal bars show the ≥2 or ≤2 cutoffs for fold change and the vertical dashed bar shows the p<0.05 cutoff for significance. Proteins highlighted in red are those with a fold change ≥ 2 and p value < 0.05 that were used for further pathway enrichment analysis. Other proteins were denoted as non-significant (ns), grey. Several significantly identified proteins are highlighted. (**B**) Pathway enrichment analysis of BioID2 significantly enriched proteins (fold change ≥ 2 and p value < 0.05) showing fold enrichment of GO terms associated with cytoskeletal elements of the sarcomere and with ribosomes and protein translation. The number of proteins identified for each term is shown as a circle and the -log10 of the false discovery rate (FDR) is color coded. (**C**) Unsupervised clustering of enriched GO terms showing identified proteins were predominantly associated with two groups – Cytoskeletal sarcomere proteins or ribosomal components. The enrichment FDR of specific GO terms is shown. (**D** and **E**) Representative immunofluorescence images (**D**) and line scan (**E**, yellow line) analysis of isolated adult rat ventricular cardiomyocytes stained for α-actinin (red), RPSA (green) and DAPI (blue) showing localization of endogenous RPSA to both sides of the Z-line identified with α-Actinin. Scale bar = 10 µm.

**Table 1.**
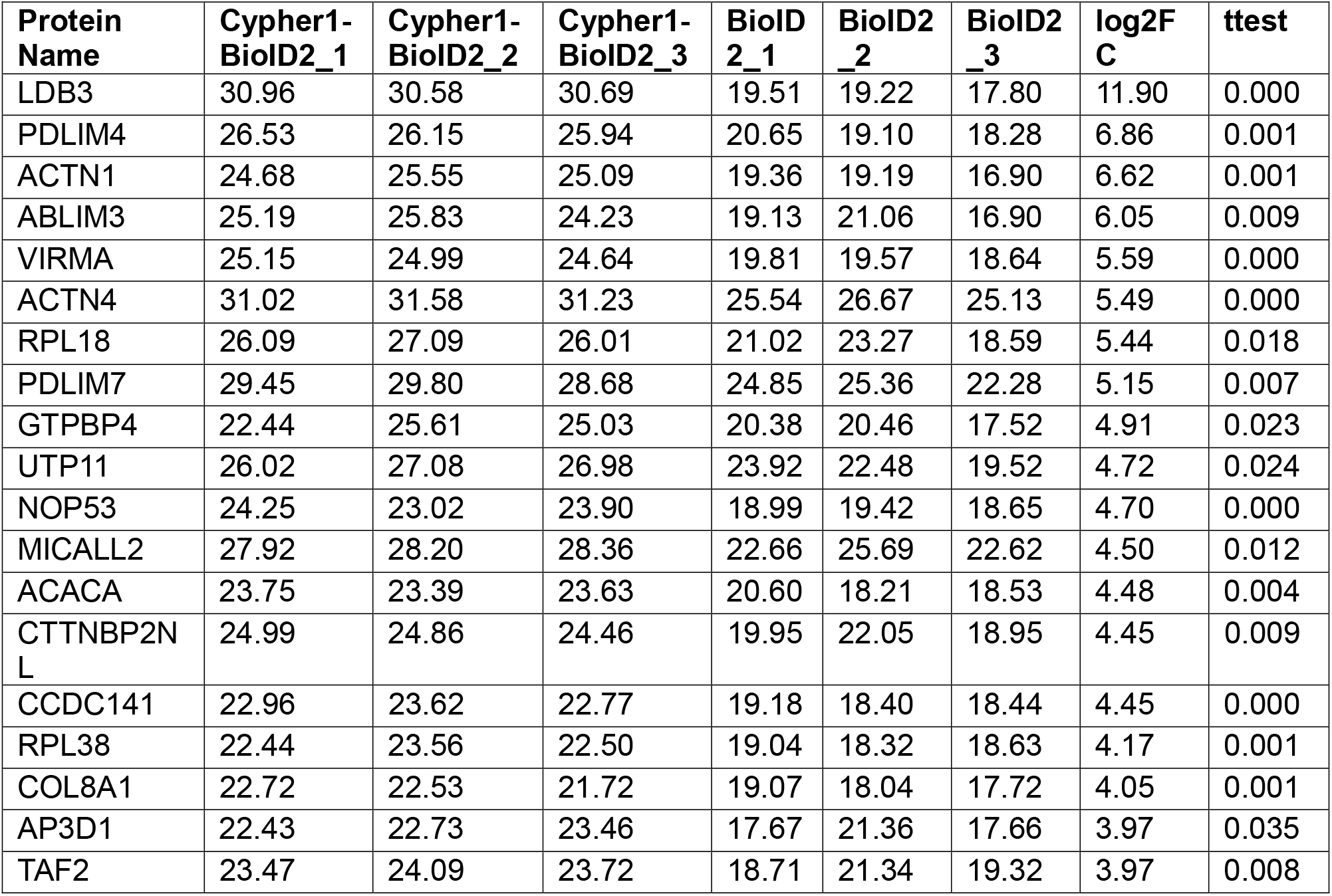
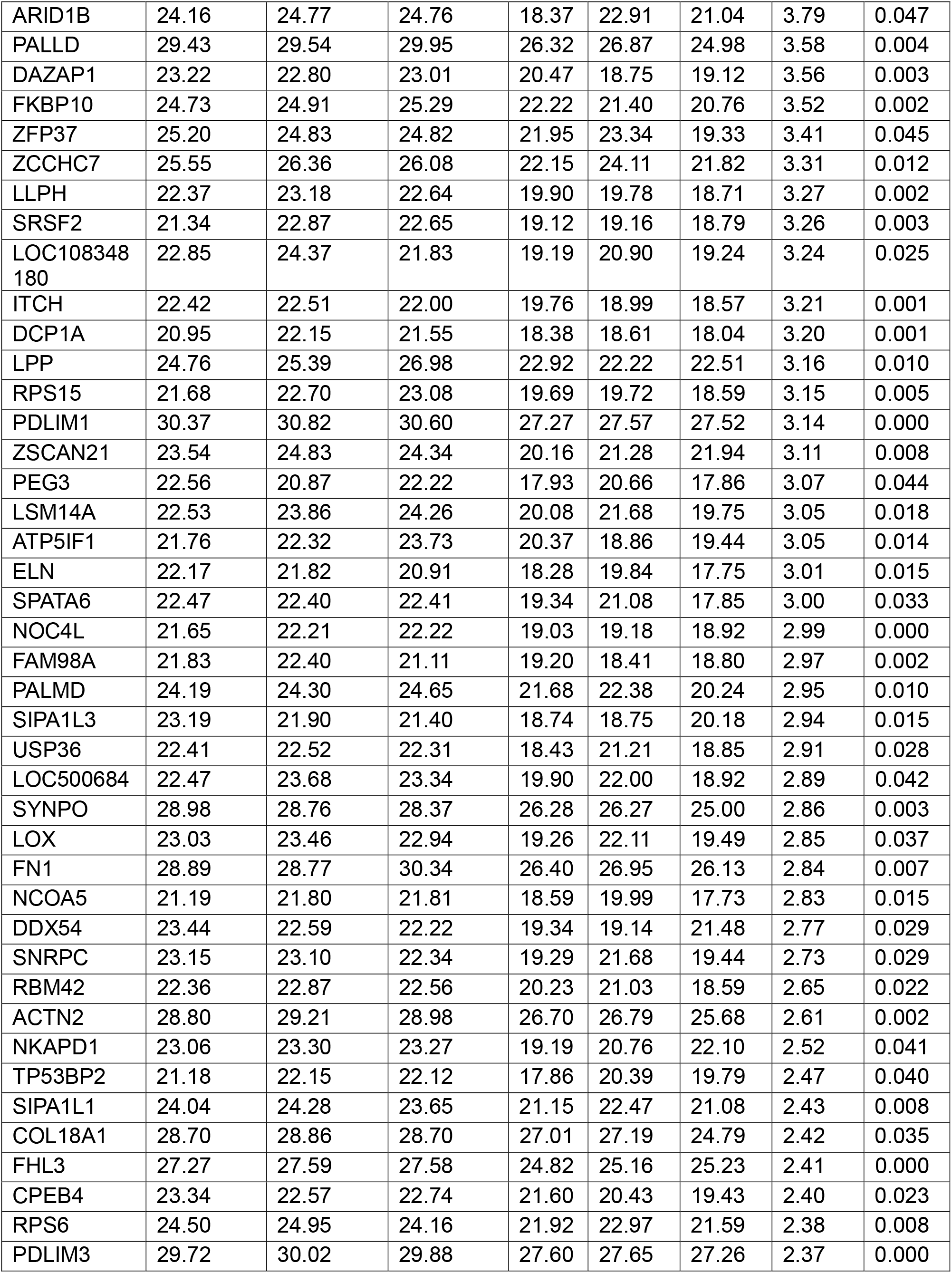
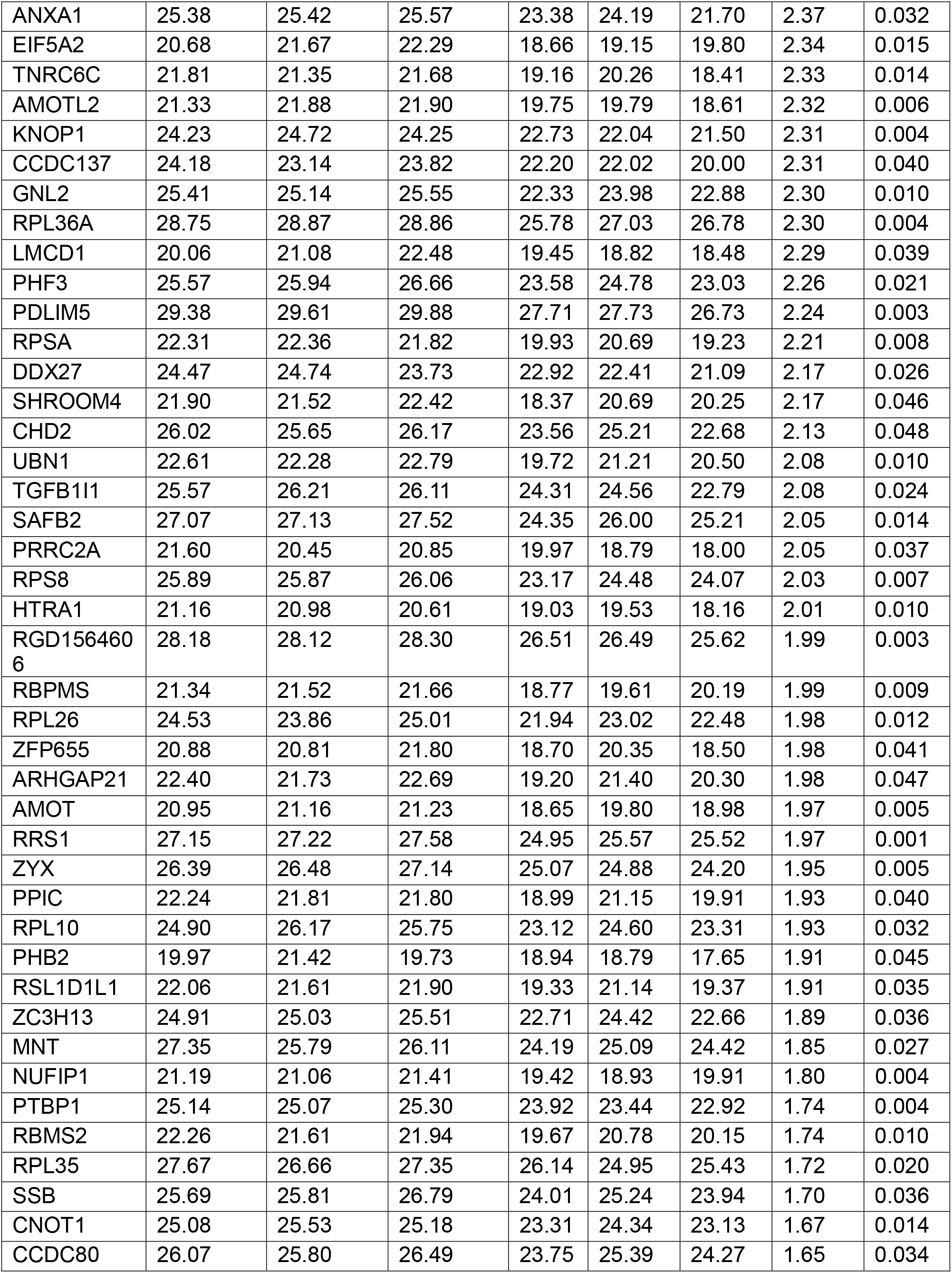
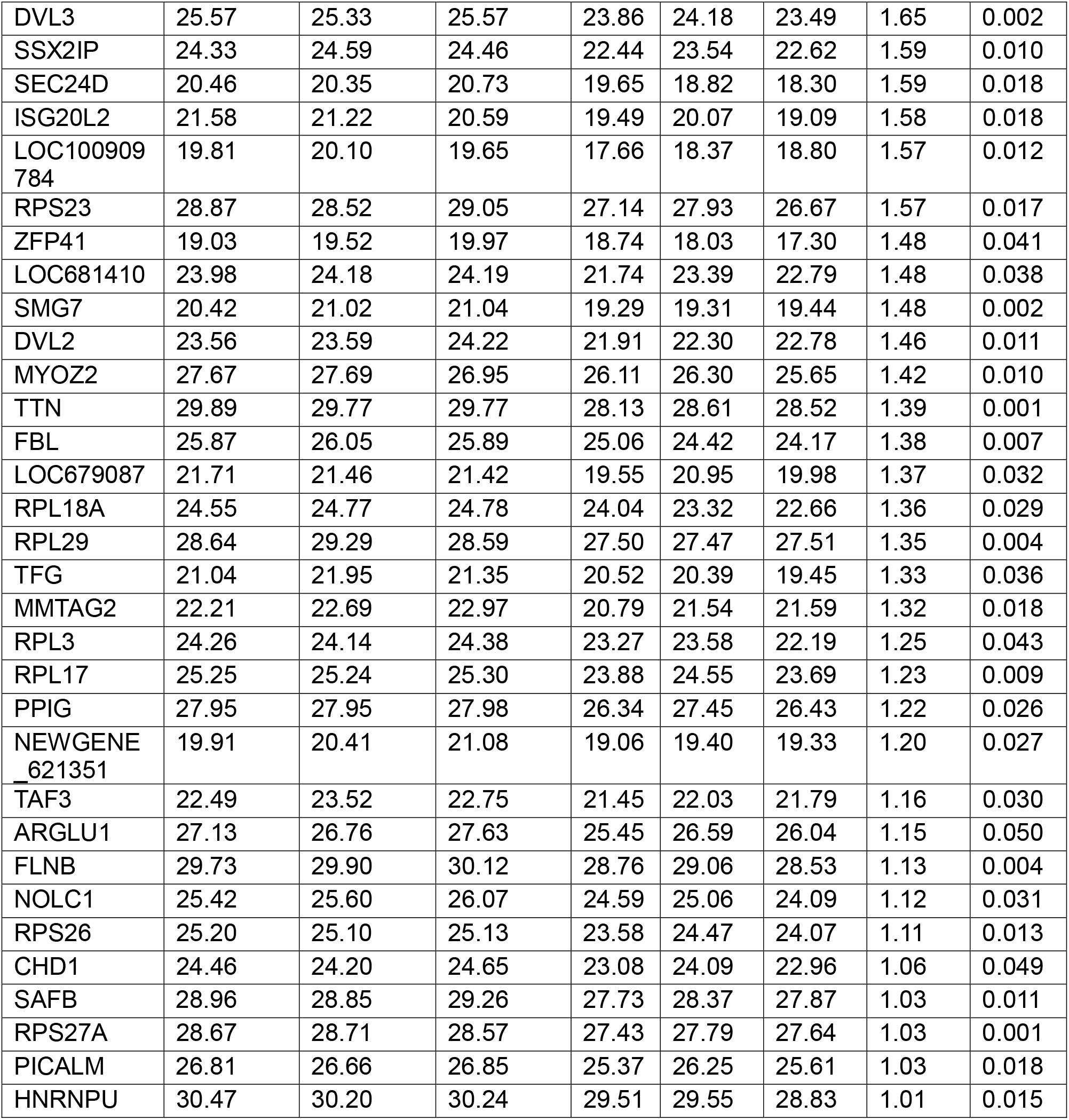
Identified proteins by Cypher-BioID2 and BioID2 triplicates. The LFQ values for each sample are shown with the Log2 of the fold change and the t-test p value.

To gain insight into the protein groups associated with the Cypher-BioID2 significantly enriched proteins (fold change ≥ 2 and p value < 0.05), we performed an enrichment analysis using gene ontology (GO) terms. The analysis indicated that the Cypher-BioID2 associated proteins were significantly enriched in several terms, including ribosome and sarcomere Z-line, further confirming our previous observation that the Z line region contains a protein translation nanodomain (Figure 2B). There is an inherent overlap among GO terms, as a given protein can be associated with multiple terms. We therefore also employed an unsupervised clustering approach with the aim of identifying distinct clusters of enriched GO terms. This analysis revealed two major clusters of protein functions. The first cluster consisted of pathways related to translational machinery, such as ribosomal proteins and ribonucleoprotein complexes. The second cluster included pathways involved in cytoskeletal elements, particularly those associated with sarcomeric structures (Figure 2C). All Cypher-BioID2 enriched proteins are also shown in a STRING analysis (Supplemental Figure 1).

Of the identified proteins we decided to concentrate on Ribosomal Protein SA (RPSA). RPSA is considered a ribosomal protein which has acquired the function of laminin receptor during evolution. Although its function in the heart is not known, it was reported to bind both ribosomes and the microtubular system (10). We also studied Ribosomal Protein L38 (RPL38), a known component of the 60S subunit of the ribosome. We performed immunofluorescence staining of RPSA in adult rat cardiomyocytes as well as three-dimensional reconstruction of Z-stacked images showing RPSA was aligned on both sides of the α-actinin containing Z lines (Figure 2, D and E, and Supplemental Video 1). This result further validates our proximity labeling approach and confirms that RPSA is localized to sarcomeric Z lines.

### Loss of RPSA disrupts localized translation and sarcomeric integrity in cultured cardiomyocytes

The localization of RPSA on both sides of the Z-line suggested that in cardiomyocytes this protein has a ribosomal rather than a laminin receptor function. We therefore asked if RPSA is required for localized protein translation in the sarcomere and for sarcomere maintenance. We used siRNA to knockdown *Rpsa* in NRVMs, or knocked down *Rpl38*, a known ribosomal protein as a control. The efficacy of the *Rpsa* knockdown was validated by quantitative PCR (RT-qPCR), which demonstrated an ∼85% reduction in *Rpsa* and ∼82% reduction in *Rpl38* mRNA levels compared to the non-targeting negative control siRNA group (Figure 3A). A reduction in RPSA protein levels was also confirmed using Western blotting (Figure 3B).

**Figure 3.**
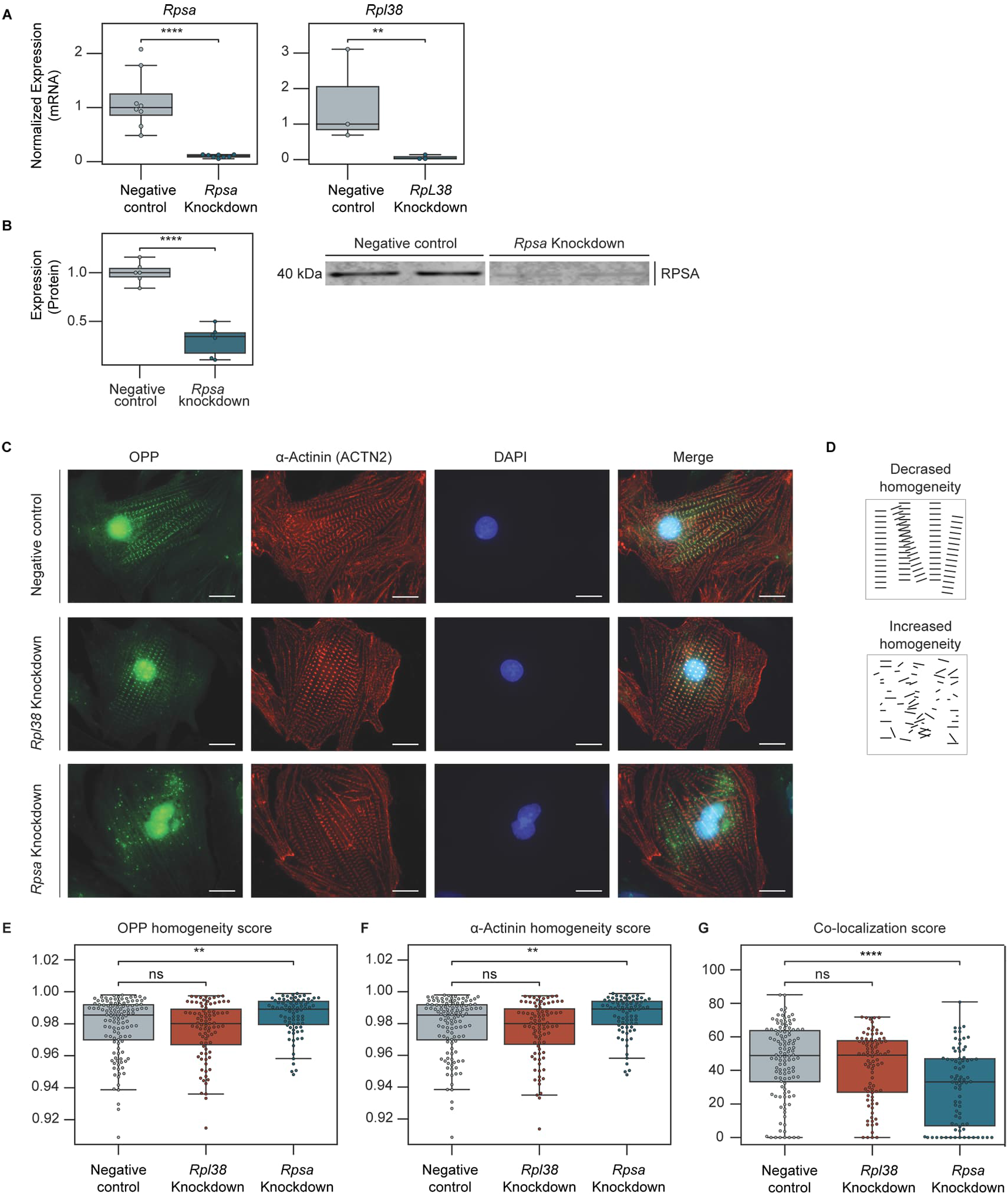
*Rpsa* Knockdown disrupts localized translation in cultured cardiomyocytes. (**A**) Gene expression analysis with RT-qPCR of *Rpsa* (left) and *Rpl38* (right) shows ∼85% and ∼82% reduction in expression levels following siRNA-mediated knockdown respectively compared with control siRNA. Expression was normalized to *Gapdh*. N = 3-8 biological samples per experimental group pooled from 3 independent experiments. (**B**) Western blot analysis (right) with densitometric analysis (left) of RPSA protein expression shows a∼70% reduction in RPSA protein level compared to control following siRNA-mediated knockdown. N = 6 biological samples per experimental group from 3 independent experiments. (**C**) NRVMs were incubated with o-propargyl-puromycin (OPP) to label nascent proteins 48 hours after treatment with negative control (top), *Rpl38* (middle), or *Rpsa* (bottom) siRNAs. Representative images of staining for OPP-labeled nascent proteins (green), α-actinin (red) and DAPI (blue) show disruption of localized protein synthesis following *Rpsa* knockdown but not following *Rpl38* knockdown or under control conditions. Scale bar = 10 µm. (**D**) Cartoon to illustrate a structure with decreased homogeneity (top, score=0.959) and a structure with increased homogeneity (bottom, score=0.977). (**E** and **F**) Homogeneity scores for translation from the OPP signal (**E**) and for the sarcomeric structure from the α-Actinin signal (**F**), showing increased homogeneity implying decreased localization following *Rpsa* knockdown but not following *Rpl38* knockdown or control siRNAs. N = 119-172 cells per group pooled from 4 independent experiments. (**G**) Co-localization analysis between the OPP and the α-Actinin signal showing significantly decreased co-localization between translation and the Z-line after knockdown of *Rpsa*. N = 119-172 cells per group pooled from 4 independent experiments. ns (p > 0.05), * (p ≤ 0.05), ** (p ≤ 0.01), *** (p ≤ 0.001), **** (p ≤ 0.0001), by unpaired Student’s t-test (A and B), 1-way ANOVA test (E-G). Data are presented as individual values with box plot displaying the median with 25th and 75^th^ percentiles.

To assess the effect of *Rpsa* or *Rpl38* knockdown on localized protein translation, the control and knockdown cells were incubated with OPP, a puromycin analogue that labels newly translated proteins, for 60 minutes. Cardiomyocytes were fixed, OPP incorporated nascent peptides were labeled using Alexa Fluor 488 by a Click reaction and immunostained with antibodies against sarcomeric ACTN2 to examine the sarcomeric structure. This analysis revealed a major disruption in localized protein synthesis with knockdown of *Rpsa*, but not with knockdown of *Rpl38* or control siRNA knockdown (Figure 3C). The sarcomere structure on the other hand appeared largely intact.

To quantify these changes, we used an automated image processing approach inspired by the Sarcomere Texture Analysis (SOTA) method (11). These custom tools segment cells from multi-channel immunofluorescence images and employ algorithms to quantify two aspects of the sarcomeric structure, namely homogeneity and co-localization of different markers.

Homogeneity score is a measure of the uniformity of signal distribution in the image, with a higher score indicating a more diffuse signal distribution, and a lower score indicating a more localized distribution (Figure 3D). This unbiased automated analysis showed that the knockdown of *Rpsa* resulted in a statistically significant increase in homogeneity in the OPP signals (Figure 3E), indicating a loss of localized protein translation with a shift toward more diffuse translation. While our visual analysis did not reveal a major disruption in the sarcomere organization, our automated analysis of the α-actinin (ACTN2) staining demonstrated an impairment (Figure 3F). The automated analysis also revealed a significant decrease in the co-localization between the ACTN2 and OPP signals after *Rpsa* knockdown (Figure 3G), indicating a much greater loss of sarcomeric localized translation relative to the changes in sarcomeric structure. In contrast with these findings, the loss of RPL38 did not affect the localization of translation or the integrity of the sarcomere. Together these data indicate that the loss of RPSA results in the loss of sarcomeric Z-line protein translation and a likely secondary loss of sarcomere structure in cardiomyocytes.

### Loss of RPSA results in cardiac disfunction in-vivo

We next wanted to study the consequences of RPSA loss in cardiomyocytes in vivo. We first used mice generated by crossing the Rosa26-Lox-stop-Lox-Cas9 knockin mice, which have a floxed-STOP cassette preventing the expression of the downstream CAS9 endonuclease from a CAG promoter knocked in the Rosa26 locus and the β-actin driven CRE recombinase strain, resulting in mice with ubiquitous and strong expression of CAS9 under the control of the CAG promoter in multiple tissues, including the heart. Studies in the Jackson Laboratory showed that constitutive CAS9 expression does not result in any detectable toxicity or morphological abnormalities (12). To achieve *Rpsa* knockout we injected neonatal (3-5 days post-birth) mice with AAVs encoding for sgRNAs targeting exons 3, 4 and 5 of the *Rpsa* gene, in equal proportions (Supplemental Figure 2A). The resulting mice are genetic mosaics in which the fraction of cardiomyocytes with *Rpsa* knockout in the heart depends on the transduction efficiency, and we refer to these mice as unconditional *Rpsa* knockout mice. For controls, we injected mice with an AAV encoding for a non-targeting sgRNA sequence. We confirmed the CRISPR-CAS9-mediated gene editing in the *Rpsa* gene in the mouse heart tissue using a T7 endonuclease I (T7E1) assay at the age of three months (Supplemental Figure 2B). We analyzed these unconditional *Rpsa* knockout mice at the age of 3 months. Echocardiography analysis was consistent with dilated cardiomyopathy with increased diastolic and systolic left ventricular dimensions, decreased wall thickness, and reduced systolic function (Figure 4, A-F). Gravimetric analysis supported these findings, showing an elevated heart weight to body weight ratio (Figure 4G).

**Figure 4.**
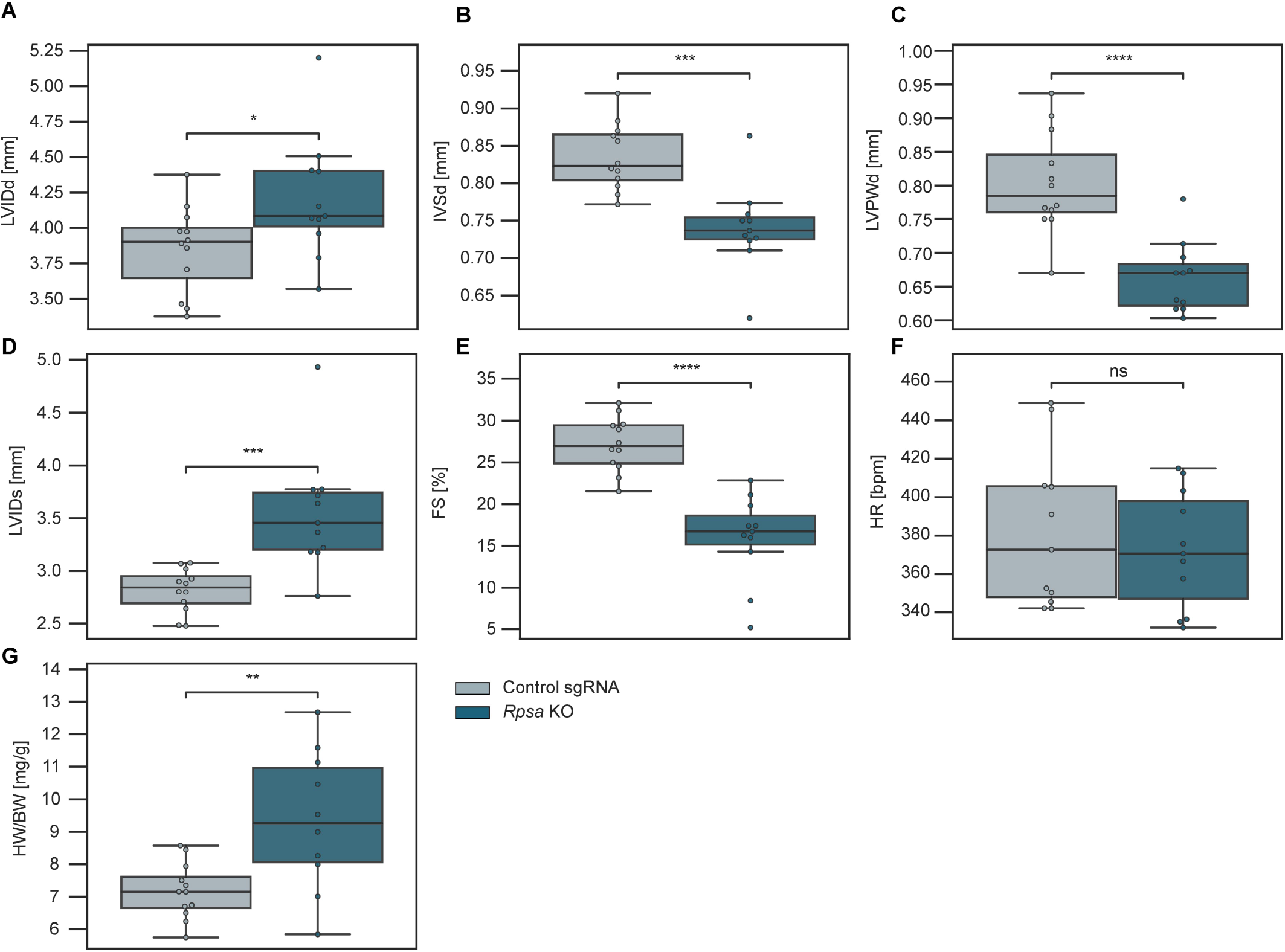
Knockout of *Rpsa* in vivo results in cardiac dysfunction. (**A**-**F**) Echocardiographic assessment of mice with unconditional CAS9 expression transduced with AAVs encoding for sgRNAs targeting *Rpsa* or control sgRNAs. Analysis shows *Rpsa* knockout results in cardiac dilation with increased left ventricular internal diameter at end diastole (LVIDd), wall thinning with decreased interventricular septal and left ventricular posterior wall diameters at end diastole (IVSd and LVPWd) and decreased contractile function with increased left ventricular internal diameter at end systole (LVIDs) and decreased fractional shortening (FS%). Heart rate (HR) was unchanged. N = 11-12 mice per group, mean of 3 measurements for each mouse. (**G**) Gravimetric analysis showing increased heart weight normalized to body weight (HW/BW) after *Rpsa* knockout. N = 11-12 mice per group. ns (p > 0.05), * (p ≤ 0.05), ** (p ≤ 0.01), *** (p ≤ 0.001), **** (p ≤ 0.0001), by unpaired Student’s t-test. Data are presented as individual values with box plot displaying the median with 25th and 75^th^ percentiles.

To ensure that the phenotype resulted from cardiomyocyte cell autonomous loss of *Rpsa* we next used the parental Rosa26-Lox-STOP-lox-Cas9 knockin mouse line. These mice were similarly transduced with AAVs that encode for the sgRNAs targeting *Rpsa*, or control sgRNA, and also express HA-tagged nuclear localized CRE under the control of the chicken cardiac troponin T promoter to delete the STOP cassette and express CAS9, specifically in cardiomyocytes. We also confirmed the CRISPR-CAS9-mediated gene editing in the *Rpsa* gene in the mouse heart tissue using a T7 endonuclease I (T7E1) assay in these conditional mice at the age of three months (Supplemental Figure 2C). We analyzed mice with cardiomyocytes specific CRE expression at the age of 1, 3, and 6 months by echocardiography (Figure 5, A-F). Interestingly, the fractional shortening was mildly increased in the *Rpsa* knockout mice at the age of 1 month. The cause of this increase is not clear and may represent a compensatory neurohumoral response to pathological changes in these hearts. By the age of 3 months there was a trend toward dilation and reduced systolic function, and the same phenotype as in the unconditional *Rpsa* targeted mice was fully manifested at the age of 6 months, with significant chamber dilation and reduced function. The delay in the development of overt cardiomyopathy in the conditional *Rpsa* knockout mice may be due to decreased efficiency of the Troponin T driven CRE expressing AAVs compared to the germline β-actin driven CRE or due to the added effect of *Rpsa* deletion in non-cardiomyocytes in the unconditional *Rpsa* knockout mice. Nevertheless, these data confirm that *Rpsa* knockout induces cardiomyocyte cell-autonomous dilated cardiomyopathy phenotype.

**Figure 5.**
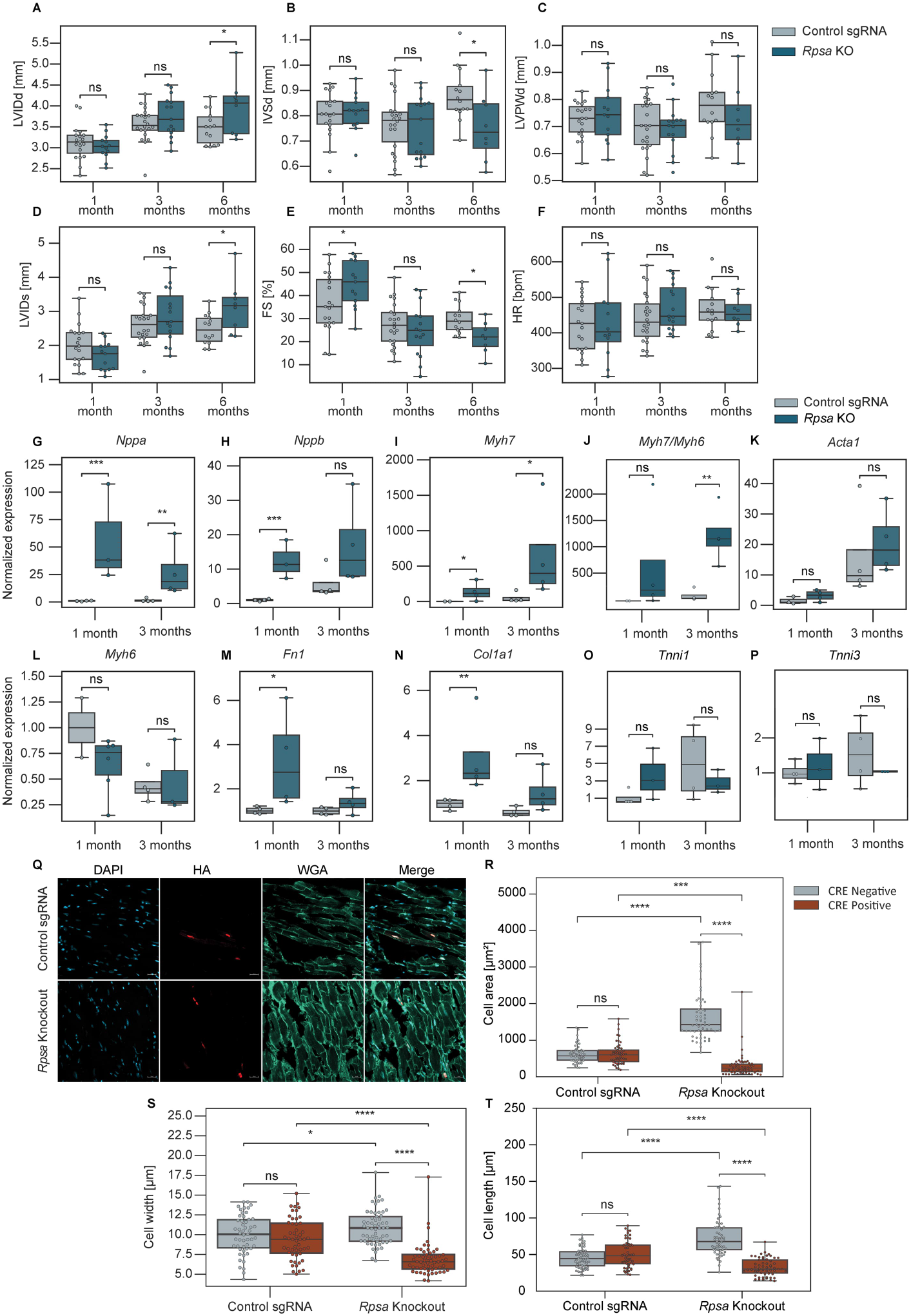
Cardiomyocyte cell autonomous loss of *Rpsa* results in cardiac dysfunction and cardiomyocyte atrophy. (**A**-**F**) Echocardiographic assessment of mice with Lox-stop-Lox-CAS9 transduced with AAVs encoding for sgRNAs targeting *Rpsa* or control sgRNAs and troponin promoter driven HA-CRE expression to knockout *Rpsa* specifically in cardiomyocytes. Analysis shows *Rpsa* knockout results in cardiac dilation with increased left ventricular internal diameter at end diastole (LVIDd), decreased interventricular septal and left ventricular posterior wall diameters at end diastole (IVSd and LVPWd), and decreased contractile function with increased left ventricular internal diameter at end systole (LVIDs) and decreased fractional shortening (FS%). Heart rate (HR) was unchanged. N = 8-22 mice per experimental group. mean of 3 measurements per mouse. (**G**-**P**) Gene expression analysis by RT-qPCR shows an increase in markers of cardiac stress and hypertrophy (*Nppa*, *Nppb*, *Myh7, Myh7/Myh6 ratio*) as well as in markers of fibrosis *Fn1* and *Col1a1* in *Rpsa* knockout hearts. *Tnni1* and *Tnni3* gene expression had no statistically significant alteration. Expression data is normalized to *Gapdh*. N = 2-4 mice per age and experimental group. 2-3 technical replicates per mouse. (**Q**-**T**) Representative images and quantitative analysis in *Rpsa* knockout hearts show decreased cardiomyocyte width, length, and area in targeted HA-CRE positive cardiomyocytes, indicating atrophy, and an increase in non-targeted HA-CRE negative cardiomyocytes, indicating hypertrophy. Sections stained with Wheat germ agglutinin (WGA) (light green), HA-tag antibody (red) to identify nuclear HA-CRE, and DAPI (blue) for nuclei. N = 51-59 cells from 2-3 mice per group. Scale bar = 20µm. ns (p > 0.05), * (p ≤ 0.05), ** (p ≤ 0.01), *** (p ≤ 0.001), **** (p ≤ 0.0001), by Student’s t-test (A-P), or 1-way ANOVA test (R-T). Data are presented as individual values with box plot displaying the median with 25th and 75^th^ percentiles.

We next assessed changes in gene expression in ventricle tissue using RT-qPCR at the age of 1 and 3 months in these mice (Figure 5, G-P). Although echocardiography did not show significant changes at the age of 1 or 3 months, the fetal genes *Nppa* and *Nppb*, encoding ANP and BNP respectively, and *Myh7* were upregulated, suggesting cardiac stress. Similarly, markers of the fibrotic reaction *Fn1* and *Col1a1* were upregulated. *Acta1* trended for upregulation and *Myh6* trended for downregulation, but these changes were not statistically significant. At the age of 3 months *Nppa* and *Myh7* remained elevated, and so was the Myh7/Myh6 ratio, but *Fn1* and *Col1a1* were no longer significantly elevated. There were no significant changes in *Tnni1* or *Tnni3* levels. These findings are consistent with dilated cardiomyopathy that started to develop at an early age.

For microscopic analysis we first stained cardiac sections from both *Rpsa* knockout and control mice with antibodies for HA-tag and wheat germ agglutinin (WGA) at the age of 6 months (Figure 5, Q-T). As the mice are genetic mosaics, only some cardiomyocytes were transduced and showed *Rpsa* knockouts, and we used the HA staining to identify the transduced cardiomyocytes through their HA-CRE nuclear signal. In control mice, HA-positive and HA-negative cells displayed no significant difference in area, width, or length. However, in *Rpsa* knockout mice, HA-positive cells were significantly smaller compared to HA-positive cells in control mice and had significantly smaller area, width, and length. On the other hand, HA-negative cells in *Rpsa* knockout mice were significantly larger in area, width, and length compared to HA-negative cells in control mice. The percentage of HA-positive cardiomyocytes in *Rpsa* knockout hearts was ∼15%, did not differ between the age of 1 and 6 months (Supplemental Figure 3, A and B), and similarly we could not find apoptotic cardiomyocytes on TUNEL staining (Supplemental Figure 3C). Together these findings show cell atrophy in *Rpsa* knockout cardiomyocytes and a compensatory hypertrophy of the untransduced cardiomyocytes in the mosaic *Rpsa* knockout hearts, without an evidence for significant cell death of *Rpsa* knockout cardiomyocytes.

### Loss of RPSA results in loss of localized translation and loss of ribosome localization in-vivo

Next, we wanted to see if loss of RPSA resulted in the loss of localized protein translation in cardiomyocytes in vivo. We intraperitoneally injected 3 months old *Rpsa* knockout and control mice with OPP (0.049 mg/g) and after 1 hour, we sacrificed the mice, harvested the hearts, and performed cryosections. The OPP labeling was performed by a Click reaction on the sections and the cells were co-stained for the HA-tag to identify the nuclear HA-CRE in the transduced cells. As we have previously shown (7), the OPP staining displays a characteristic striated pattern, reflecting localized sarcomeric protein translation in wild-type cardiomyocytes. When compared with HA-positive cardiomyocytes in control mice, *Rpsa* knockout cardiomyocytes lacked this localized pattern indicating that the loss of RPSA significantly reduced sarcomeric localized translation (Figure 6, A and B). The cross-striated pattern of the OPP signal was quantified by measuring the mean amplitude of a line-scan across cardiomyocytes’ long-axis, and *Rpsa* knockout cardiomyocytes showed significantly lower amplitude indicating a more diffuse signal (Figure 6C). Quantification of the density of the OPP signal showed that while translation activity was reduced in *Rpsa* knockout cardiomyocytes as compared with cardiomyocytes transduced with control sgRNA, the OPP signal was still significantly higher in *Rpsa* knockout cardiomyocytes than in mice with click reactions but without OPP (NC.1) or mice with OPP but no click reaction (NC.2), (Figure 6D). These analyses indicate that loss of RPSA results in disruption of localized translation at the sarcomeres but not a complete abrogation of protein translation.

**Figure 6.**
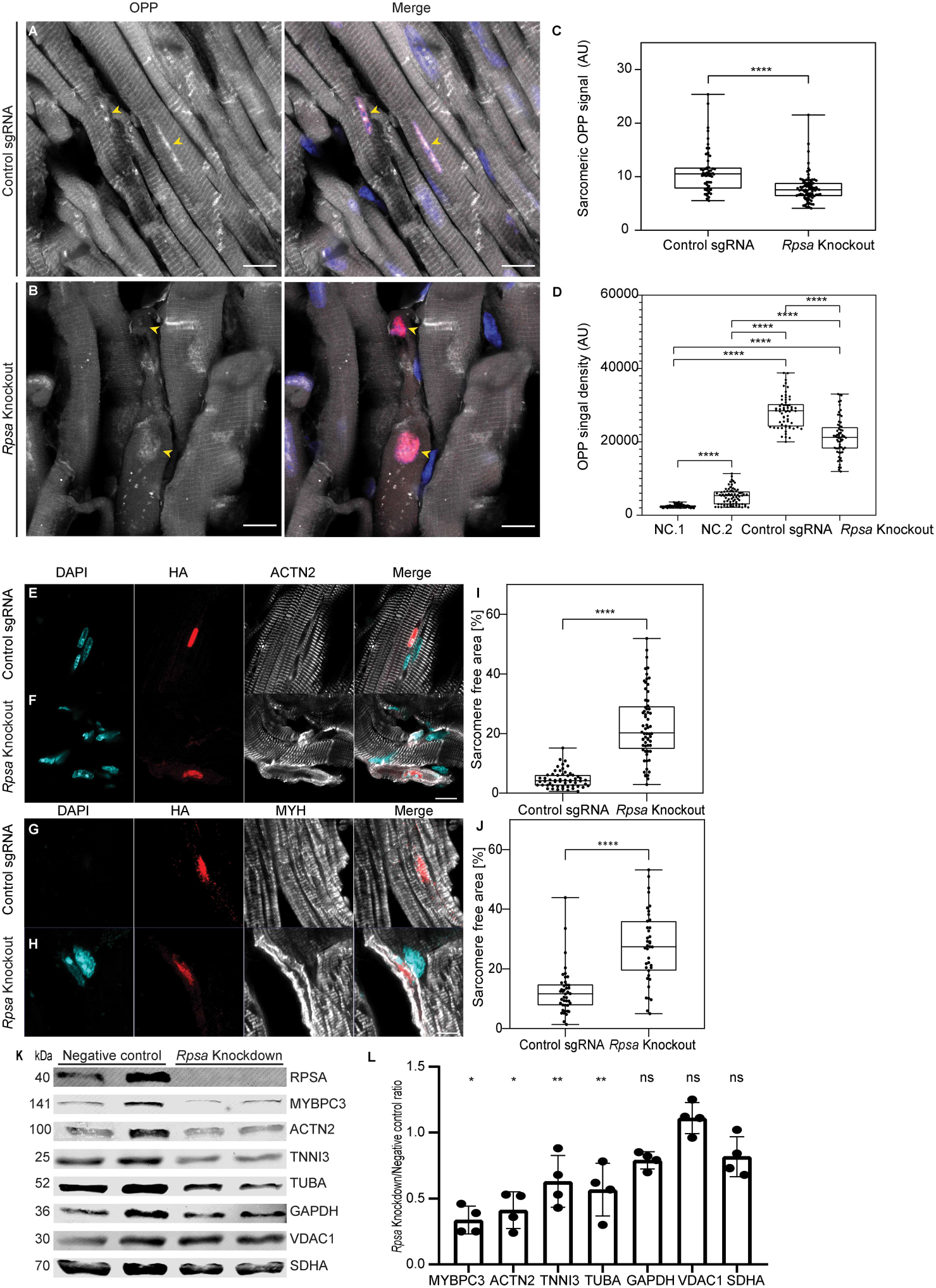
*Rpsa* knockout disrupts local sarcomeric protein translation and sarcomere integrity in-vivo. (**A**-**D**) Mosaic *Rpsa* knockout and control mice were injected with OPP and stained for OPP and HA-CRE to label nascent proteins and targeted cardiomyocytes, respectively. Representative images show OPP localized sarcomeric pattern of translation in negative control mice (**A**). In *Rpsa* knockout mice the localized sarcomeric translation is lost only in targeted HA-CRE positive cardiomyocytes (**B**). OPP-labeled nascent proteins (white), HA-tag (red) and DAPI (blue). Yellow arrowheads mark targeted nuclei. Scale bar = 10 µm. (**C**) Line-scan quantification in targeted cardiomyocytes confirms the loss of localized sarcomeric translation in *Rpsa* knockout cardiomyocytes. (**D**) The translational activity in *Rpsa* knockout cardiomyocytes, assessed by OPP cytoplasmic integrated density, is reduced yet higher than the negative control samples (NC.1-no OPP or NC.2-no Click labeling). N = 2 mice per experimental group. n = 57-88 cardiomyocytes. (**E**-**J**) Representative images and analysis showing disrupted ACTN2 and myosin heavy chain (MYH) staining in *Rpsa* knockout cardiomyocytes, quantified by the increased percentage of the sarcomere free area. N = 2 mice for Control sgRNA, N = 3 for *Rpsa* knockout group, n = 56-64 (I), n = 39-42 (**J**) cardiomyocytes. (**K** and **L**) Western blot and densitometry analysis after *Rpsa* siRNA mediated knockdown in NRVMs showing significantly reduced sarcomeric proteins levels (MYBPC3, ACTN2, TNNI3) and α-Tubulin (TUBA) but not in the non-sarcomeric proteins (GAPDH, VDAC1, SDHA). n = 4 biological samples from 3 independent experiments. ns (p > 0.05), * (p ≤ 0.05), ** (p ≤ 0.01), *** (p ≤ 0.001), **** (p ≤ 0.0001), by 1-way ANOVA test (**D**), or Student’s t-test (**C**, **I**, **J**, and **L**). Data are presented as individual values with box plot displaying the median with 25th and 75^th^ percentiles. Bar graph is shown as mean ±SD.

The loss of sarcomeric localized translation is expected to result in the loss of sarcomeric proteins. To assess this loss, we immunostained sections with antibodies to ACTN2 or myosin heavy chain (MYH), together with HA-tag to identify transduced cardiomyocytes. This analysis showed that the cytoplasm of *Rpsa* knockout cardiomyocytes largely lacked these sarcomeric proteins, with some remaining signal in the cell periphery (Figure 6, E-J). To further confirm and quantify the loss of sarcomeric proteins we knocked down *Rpsa* expression in cultured NRVMs with *Rpsa* or control siRNAs. While the knockdown of *Rpsa* resulted in a significant loss of the sarcomeric proteins MYBPC3, ACTN2, and TNNI3, or the non-sarcomeric protein tubulin, the levels of the mitochondrial proteins VDAC1 or SDHA remained mostly unchanged (Figure 6, K and L).

To verify that RPSA was indeed associated with the ribosomal complex we performed polysome profiling from normal mice heart extracts. Western blot analysis from these fractions showed RPSA was associated with both monosomes (80S) and polysomes (Supplemental Figure 4A). The association of RPSA with polysomes indicates it is part of the ribosome complex actively engaged in mRNA translation. Co-staining of sections from control mice administered OPP for RPSA showed RPSA was localized on both sides of the OPP incorporated nascent proteins at the sarcomere (Supplemental Figure 4, B and C), further supporting the association of RPSA with active translation adjacent to the sarcomere Z-line.

We next asked if the loss of localized translation resulted from impaired ribosome localization in *Rpsa* knockout cardiomyocytes. We used dual single-molecule fluorescent in situ hybridization (smFISH) with probes directed at the ribosomal 18S RNA to identify ribosomes and probes targeting the *Cre* mRNA sequence to identify transduced cells in 6-months-old mice. While HA-CRE signal was nuclear due to the nuclear localization signal in CRE proteins, the *Cre* mRNA signal was expected to be both nuclear and cytoplasmic and showed a perinuclear accumulation. In both control and *Rpsa* knockout mice, *Cre*-negative cells maintained a typical striated pattern when probed for 18S rRNA, indicative of normal ribosomal localization (8) (Figure 7, A-E). However, in *Rpsa* knockout mice, *Cre*-positive cells exhibited a loss of this striation pattern, showing a disruption in the normal sarcomeric localization of ribosomes. RPSA loss disrupted ribosome localization but did not eliminate ribosomes, explaining why localized translation was lost in the OPP assay while protein translation was not completely abrogated.

**Figure 7.**
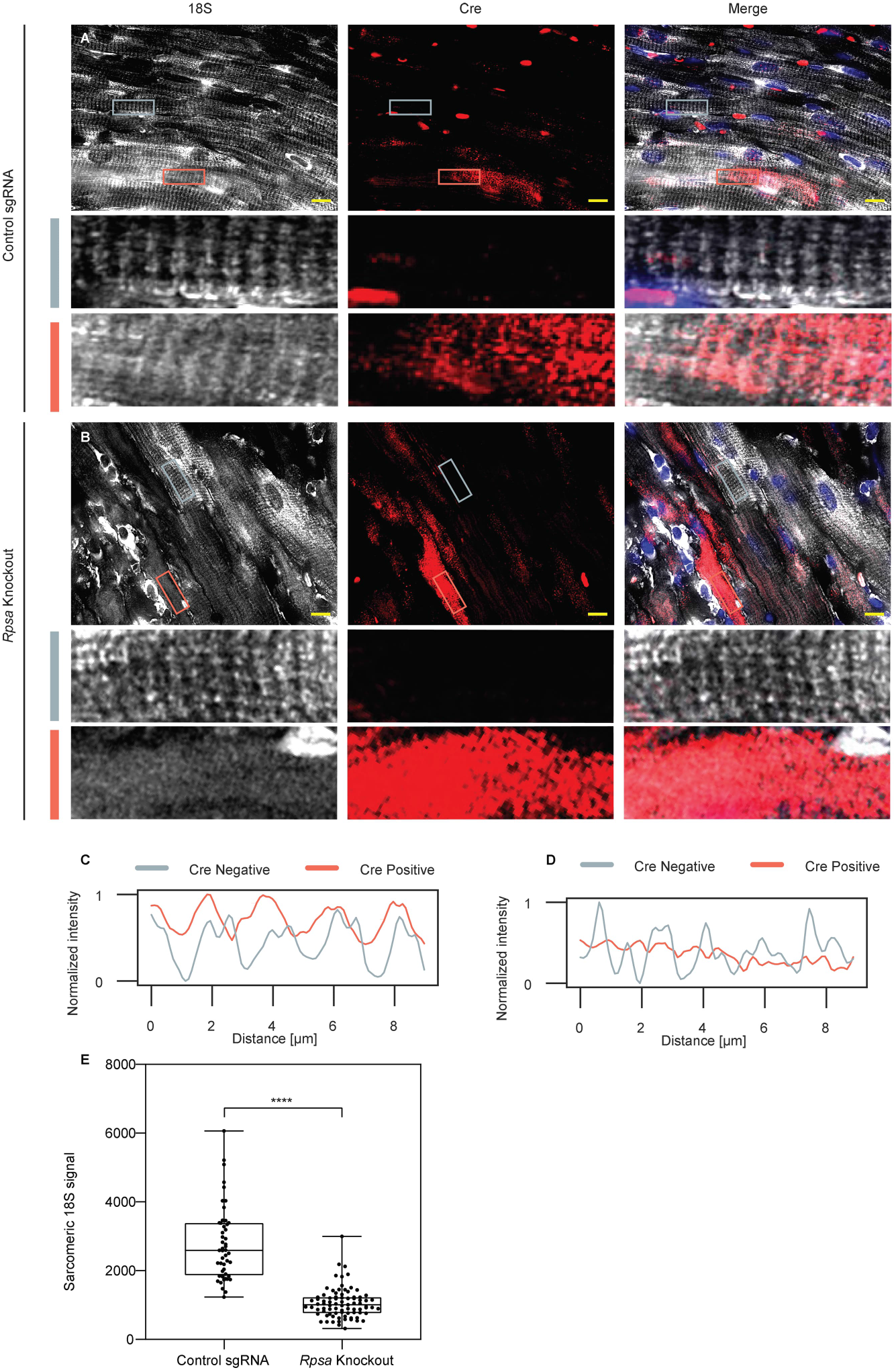
*Rpsa* knockout disrupts sarcomeric ribosome localization in-vivo. (**A** and **B**) Dual smFISH with probes for ribosomal18S RNA (white) and *Cre* mRNA (red) was used to visualize ribosomes and targeted cardiomyocytes respectively. Representative images show sarcomeric cross striated pattern of ribosomes localization in both targeted and untargeted cardiomyocytes in negative control mice (**A**). In *Rpsa* knockout mice the sarcomeric localization of ribosomes is lost in targeted *Cre* positive cardiomyocytes, while untargeted cells are normal (**B**). Inset shows magnification of untargeted cytoplasmic *Cre* mRNA negative cell (grey rectangle) and targeted cytoplasmic *Cre* mRNA positive cell (light red rectangle) for both the negative control (**A**) and *Rpsa* knockout (**B**) hearts. Scale bar = 10 µm. (**C** and **D**) The intensity of ribosomal 18S RNA FISH signal along the cardiomyocytes in the inset shows sarcomeric periodicity in both the targeted (light red line) and untargeted (grey line) cardiomyocytes in negative control mice (**C**). In *Rpsa* knockout mice (**D**) the sarcomeric periodicity of ribosomes is lost in the targeted *Cre* positive cardiomyocyte (light red line), while the untargeted cell has normal periodicity (grey line). (**E**) Line-scan quantification confirms the loss of localized sarcomeric 18S signal in *Rpsa* knockout cardiomyocytes. N = 2 mice per experimental group, n = 49-83 cells. **** (p ≤ 0.0001), by Student’s t test. Data are presented as individual values with box plot displaying the median with 25th and 75^th^ percentiles.

We also examined mRNA localization using dual smFISH with probes directed at polyA to identify mRNAs and probes targeting the *Cre* mRNA (Supplemental Figure 5). In control cardiomyocytes mRNAs displayed the expected cross striated distribution (7), (Supplemental Figure 5). Due to the small size of the *Rpsa* knockout cardiomyocytes we could not reliably quantify and determine if the mRNA distribution in these cells was intact but in some knockout cardiomyocytes we could observe a cross striated pattern.

The N-terminal domain (amino acids 1–209) of RPSA is globular highly conserved region which shares high resemblance to the prokaryotic ribosomal proteins S2 (RPS2) family (10). This region was shown to be sufficient for ribosomal binding. The C-terminal domain (amino acids 210-295) is absent in prokaryotes and has an intrinsically disordered structure. We therefore wanted to verify if the C-terminal domain of RPSA was sufficient for localization to the sarcomere. We expressed HA-tagged-mCherry-RPSA-C-Terminal fragment_210-295_ in mice using AAVs. A staining of cardiac sections from these mice with anti-HA and anti-ACTN2 antibodies showed the C-terminus of RPSA was sufficient to localize the construct at the same location as endogenous RPSA and ribosomes, on both sides of the Z-line (Supplemental Figure 6, A and B). A biochemical study showed that RPSA directly binds tubulin with high affinity (13), suggesting this interaction was mediated by the C-terminal domain of RPSA. Our protein quantification showed that knockdown of *Rpsa* resulted in reduced tubulin levels in cardiomyocytes (Figure 6, K and L), but staining of microtubules in *Rpsa* knockdown cardiomyocytes, showed that although the levels of microtubules appeared lower, the overall distribution patterns did not change (Supplemental Figure 6C). In contrast, the disruption of the microtubular network in cultured ARVMs by either colchicine or nocodazole resulted in loss of RPSA sarcomeric localization and perinuclear accumulation of RPSA (Supplemental Figure 6, D-F). These observations support the role of microtubules in RPSA localization to the sarcomere.

Last we examined the distribution and expression of RPSA in adult hypertrophic *Mybpc3_c.2373InsG_*mice, which are homozygous for the hypertrophic cardiomyopathy Dutch founder mutation (14). Immunostaining showed that RPSA was positioned on both sides of the Z-line in both hypertrophic and in wildtype cardiomyocytes (Supplemental Figure 7, A and B). Expression analysis on the other hand showed a strong trend towards downregulation of *Rpsa* in the hypertrophic hearts (Supplemental Figure 7C).

## Discussion

Here we used proximity ligation to unravel the Z-line proteome and identified RPSA as a potential localization candidate. We show that RPSA is required for localization of ribosomes, for the localization of translation, and for sarcomere maintenance in cardiomyocytes in vivo. The loss of RPSA causes the ribosomes and translation to lose their sarcomeric localization, eventually leading to dilated cardiomyopathy. In genetic mosaic mice the untraduced cells undergo compensatory hypertrophy in response to the atrophy of the *Rpsa* knockout cardiomyocytes. RPSA is known to bind the microtubular system (13), and we also show here that the C-terminus extra-ribosomal domain of RPSA is sufficient for localization to the Z-line of the sarcomere, and that disruption of the microtubules results in RPSA collapse around the nucleus.

We used Cypher (LDB3) as our bait for the spatial proteomic analysis. This protein interacts through its N-terminal PDZ domain with the Z-line protein α-actinin (15), and BioID2 was tethered to its C-terminal part. We showed that the bait is localized to the Z-line, and our proteomic analysis identified multiple know Z-line proteins including α-actinin, other PDZ-LIM proteins such as Enigma (Pdlim7) and Enigma Homologue (Pdlim5), Palladin, and Myozenin, confirming that we specifically enriched for Z-line proteins. We also identified multiple proteins associated with ribosomes and protein translation, providing further evidence that this region is an important protein translation hub. A previous study used proximity labeling experiment to unravel the Z-line proteome (16). As compared with our study, this study used human iPS-derived cardiomyocytes, which are less structurally mature than neonatal rat cardiomyocytes, a different bait, α-actinin instead of Cypher, and a different control – no BioID as opposed to untethered BioID here. While several ribosomal proteins were identified in this study, RPSA was not among the significant hits, and interestingly IGF2BP2, a major hit, was not identified in our study. The width of the Z-line measured by electron microscopy is 130 nm (17), and the exact location of the C-terminus of Cypher within the structure is not known. Given the uncertainty regarding the exact localization of Cypher C-terminus and that BioID has a practical labeling radius estimated at ∼10 nm (18), it is difficult to determine precisely where the ribosomal and translation nanodomain resides inside or near the Z-line. Our co-immunostaining of RPSA and α-actinin suggests that ribosomes or at least RPSA mostly reside on both sides of the Z-line at the lateral ends of the I-band. We speculate therefore that we mostly isolated proteins from a nanodomain located on both sides of the of the Z-line while the α-actinin-BioID study identified a domain that is more at the center of the of the Z-line.

There is controversy over the location and functions of RPSA. RPSA was separately identified as a non-integrin laminin receptor with a 67-kDa molecular weight from muscle cell membranes (19) and as a component of the 40S ribosome with a 32-37-kDa molecular weight (20) and was called 37/67-kDa laminin receptor (LAMR). It was reported to reside in the plasma membrane, the ribosomes, the cytosol, and the nucleus (10). There are many unanswered questions regarding the cellular biology of this protein, about its ability to bind laminin, and about the function of its intrinsically disordered C-terminal region (10). While RPSA’s N-terminal domain (amino acids 1-209) is homologous to the bacterial small ribosomal protein rps2, its C-terminal domain (amino acids 210-295) is not found in prokaryotes. It was therefore proposed that the N-terminal domain binds ribosomes while the C-terminal domain of RPSA functions in laminin binding. Nevertheless, it was also proposed that the C-terminal domain of RPSA participates in ribosome binding (21).

It is also unclear how RPSA functions as a ribosomal protein. It is highly abundant, with six to eight copies per ribosome, and does not appear to be a constitutive component of ribosomes (21–23). We found that in cardiomyocytes RPSA is localized to both sides of the Z-line and near the intercalated discs. This location coincides with our previously documented location of ribosomes (7, 8), and together with the proteomic analysis here, that identified other ribosomal protein at this site, suggests that RPSA is predominantly associated with ribosomes near the Z-lines of cardiomyocytes. Our polysome profiling showed RPSA was associated with monosomes and polysomes, but it is unclear whether or not all cardiomyocytes’ ribosomes contain RPSA or use it for localization. We showed that the sarcomeric localization of ribosomes was lost after deletion of RPSA in vivo. In cardiomyocytes and muscle cells, the bulk of the transcriptome is located along the Z-line (7, 8), including transcripts for non-sarcomeric proteins (9). We examined mRNA localization in *Rpsa* knockout cardiomyocytes using dual smFISH for polyA. The small size and relatively weak signal of dual color FISH made it difficult to determine whether the cross striated localization of mRNA was still intact in these knockout cardiomyocytes. However, we and others have previously shown that the sarcomeric mRNA distribution does not depend on ribosomes and translation (7, 9). The untethering of ribosomes from the Z-line should increase the distance between ribosomes and mRNAs, resulting in a decrease in protein translation. Indeed, we show that localized translation in cardiomyocytes was lost after knockdown of *Rpsa* in cultured cardiomyocytes or by CRISPR knockout in vivo with a loss of sarcomeric proteins.

Although knockout of *Rpsa* resulted in cardiomyocyte atrophy, we could identify knockout cardiomyocytes even several months after RPSA ablation and could show these cardiomyocytes contained ribosomes and exhibited translation, albeit reduced and unlocalized. We also did not observe a significant decline in the numbers of knockout cardiomyocytes between 1 and 6 months of age or evidence for apoptosis. Considering the limited half-lives of proteins in the heart (24), these results strongly suggest that RPSA is not required for ribosome biogenesis or protein translation in general. The loss of sarcomeric but not of mitochondrial proteins after *Rpsa* knockdown also supports this view. However, we cannot rule out that RPSA has additional ribosomal functions in addition to localization. This question is further complicated by the fact that ribosomes are responsible for the translation of ribosomal proteins, and therefore the untethering of ribosomes from the Z-lines and from the localized mRNAs there is also expected to reduce the translation of ribosomal proteins, leading to further reduction in translation. Our data also suggests that the atrophy and functional loss of ∼15% of cardiomyocytes is sufficient for the development of heart failure.

Proteomic pull-down in fibroblast showed that RPSA interacts with cytoskeletal elements in addition to its interactions with multiple ribosomal proteins (25). In particular, elements of the microtubular system including tubulin and Dynein cytoplasmic 1 intermediate chain 2, a molecular motor protein responsible for transport of proteins along microtubules were identified.

A biochemical study showed that recombinant RPSA, but not the bacterial orthologue rps2, directly bind both tubulin and actin with high affinity, suggesting that the C-terminal domain of RPSA, that is absent in bacteria, mediates this interaction (13). This study also suggested that ablation of the microtubular system results in altered distribution of ribosomes in cultured fibroblasts. We have previously shown that the sarcomeric Z-line association of ribosomes and translation is lost upon ablation of the microtubular system (7, 8). Together with our observations that the C-terminal domain of RPSA is sufficient for binding to the Z-line, and that RPSA loses its sarcomeric localization after disruption of the microtubular system these data imply that RPSA ability to bind to the microtubular system by its C-terminal domain and its ability to bind ribosomes is responsible for ribosomal Z-line localization of ribosomes and translation in cardiomyocytes. We cannot rule out however that the localization is mediated by binding of RPSA to additional proteins.

A spontaneously occurring mutation in *Rpsa* in a mouse line resulted in arrhythmogenic right ventricular cardiomyopathy, that was ascribed to nuclear functions of RPSA and binding to heterochromatin protein 1, suggesting that this gene may have a specific function in the heart (26). We show here that mice with the homozygous Dutch *Mybpc3* mutation, which results in complete loss of MYBPC3 have a strong trend for reduced expression of *Rpsa*. Further research is needed to determine how such cardiac disease associated *Rpsa* reductions contribute to disease progression. The homozygous *Rpsa* knockout mice die at an early embryonic age, and therefore the necessity for this gene could not be studied in vivo (10). In this study, we used a cardiomyocyte-specific genetic mosaic knockout of *Rpsa* to investigate the cardiomyocyte cell-autonomous effects of RPSA loss in post-natal cells within the context of normally functioning myocardium, although the mice eventually succumbed to cardiomyopathy. A knockout of *Rpsa* in all cardiomyocytes would have likely resulted in profound cardiac dysfunction and would have made the dissection of the direct effects of RPSA deletion the from secondary effects difficult (27).

The role of RPSA in tethering ribosomes to microtubules has been proposed before (13), but the highly localized and cross striated distribution of ribosomes and translation in cardiomyocytes allowed us to examine and demonstrate the necessity of RPSA for localized translation in vivo. It is likely however that RPSA is required for localized translation in many cell types. Multiple mechanisms were shown to be responsible for mRNA localization in different cell types and cellular locations (28), and therefore it is possible, or even likely, that additional mechanisms and elements are responsible for ribosome and translation localization beyond RPSA in muscle cell and in additional cell types.

## Methods

### Sex as a biological variable

Our study examined male and female animals equally, and similar findings are reported for both sexes.

### sgRNAs

(see table 2)

**Table 2.**
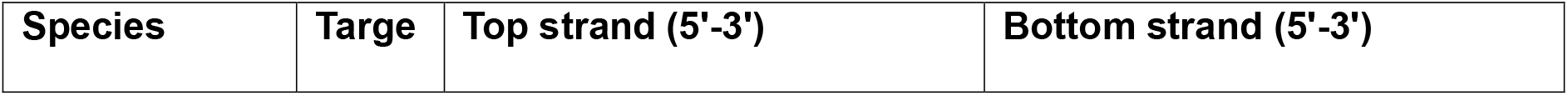
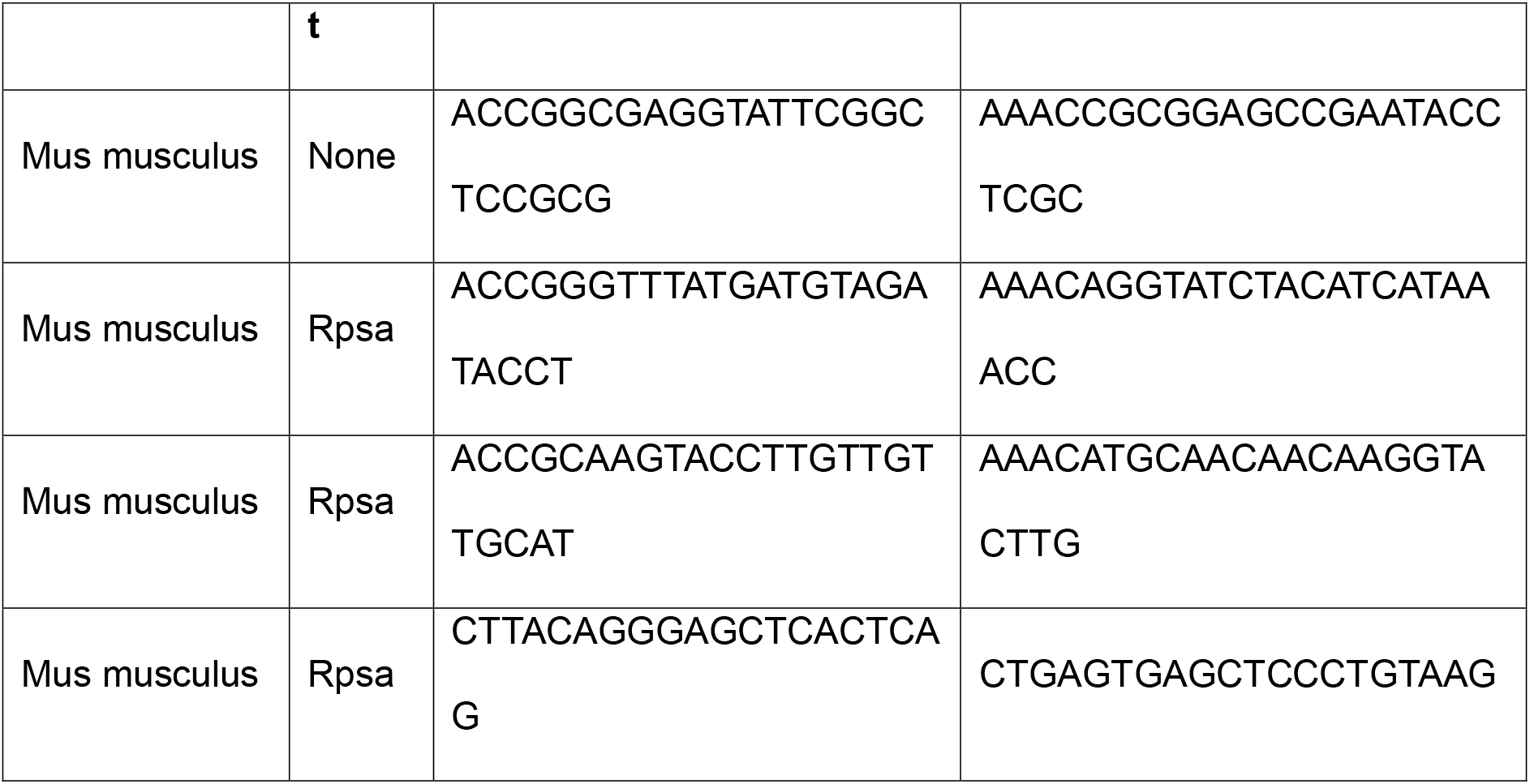
sgRNAs.

### Antibodies

(see table 3)

**Table 3.** Antibodies.

| **Name** | **Manufacturer** | **Catalog #** |
| --- | --- | --- |
| Anti-Cardiac Troponin I antibody | Abcam | ab47003 |
| Anti-alpha-Actinin antibody | Sigma-Aldrich | A7811 |
| Anti-BioID2 antibody [SS 3A5-E2] | Abcam | ab232733 |
| Anti-67kDa Laminin Receptor antibody [EPR8469] | Abcam | ab133645 |
| Anti HA-tag antibody | Abcam | ab9110 |
| Anti MYBPC3 antibody | Abcam | Ab262964 |
| Anti RPS6 antibody | Cell Signaling Technology | C-2217 |
| Anti RPL10A antibody | Abcam | ab174318 |
| Anti-alpha TUBULIN antibody | Cell Signaling Technology | 3873S |
| Anti-MYH antibody | DSHB | A4.1025 |
| Donkey Anti-Rabbit IgG antibody IRDye® 680RD | Li-Core Bioscience | 926-68073 |
| Goat Anti-Mouse IgG Antibody, IRDye® 800CW | Li-Cor Bioscience | 926-32210 |
| Lectin from Triticum vulgaris (wheat), FITC conjugate | Sigma Aldrich | L4895 |
| Click-iT TUNEL Alexa Fluor 488 Imaging Assay | Thermo-Fisher Scientific | C10245 |

### Plasmids

pAAV2/9n (Addgene plasmid #112865; http://n2t.net/addgene:112865; RRID: Addgene_112865) and pAdDeltaF6 (Addgene plasmid #112867; http://n2t.net/addgene:112867; RRID: Addgene_112867) were a gift from James M. Wilson. mycBioID2-pBABE-puro was a gift from Kyle Roux (Addgene plasmid #80900; http://n2t.net/addgene:80900; RRID: Addgene_80900).

### Isolation of cardiomyocytes and tissue culture

NRVMs were isolated using the Worthington Biochemical Neonatal Cardiomyocyte Isolation system (Cat. #LK003300) according to the manufacturer’s instructions. Cardiomyocytes were separated using Percoll gradients. Cytosine β-D-arabinofuranoside (ARA-C, Sigma-Aldrich #C1768) was added to the culture medium for a final concentration of 3 µM to inhibit the proliferation of fibroblasts. Isolation of ARVMs was performed according to the previously published protocol (29).

### Production of adenoviruses and adeno-associated Viruses

Adenoviruses were generated as previously described with adenovirus expression plasmid containing the insert (Chypher-BioID2 or BioID2) (7). The production of AAV followed a previously published protocol (30). HEK293 cells were transfected with the pAAV plasmid, along with a helper plasmid (pAdDeltaF6) and a packaging plasmid (pAAV2/9n). For injection of AAV to mice, 50 µL of containing approximately 3*10^11^ viral genomes were injected into the intraperitoneal cavity of 3-5 days old mouse pups.

### Microscopy

Samples were imaged using a Zeiss LSM 900 with Airyscan2 super-resolution system, attached to an Axio Observer 7 inverted microscope. Acquisition was performed with Plan-Apochromat 63x/1.4 oil DIC M27 objective from Zeiss. Airyscan images were processed with the ZEN software using default parameters.

For widefield imaging, slides were imaged with Axio Observer inverted fluorescent microscope (Zeiss) using an X-cite metal-halide light source and a high-resolution camera (Hamamatsu Orca R2), with an X63/1.4NA objective (Olympus).

### Microtubule Ablation

The cell culture medium was replaced with a medium containing either Nocodazole or Colchicine at a final concentration of 10µM. After a 24-hour incubation period, the cells were fixed and stained.

### Image analysis

For automated image analysis of OPP and immunofluorescence signals in NRVMs a custom tool based on the Sarcomere Texture Analysis (SOTA) method (31) was developed. Briefly, cells were segmented from multi-channel immunofluorescence images and algorithms were used to quantify two aspects of the sarcomeric structure, namely homogeneity and co-localization of different markers. To quantify the sarcomeric signal of OPP or 18S in sections, lines were drawn using Fiji software along the long axis of cardiomyocytes. We measured the mean peak-to-valley intensity along the line. We used two lines per cell and the average value from these two lines was used as the value for the cell. To quantify the signal density, we used two different regions of interest (ROI) of fixed size in the cytoplasm and calculated the integrated density of signal using Fiji. The average value from these regions was used as the value for the cell. To quantify the sarcomere free area, we manually measured and divided the area devoid of sarcomeric signal in the cytoplasm by the total area of the stained cardiomyocyte in the sections using Fiji.

### Immunofluorescence staining

Immunofluorescence staining was performed on isolated cardiomyocytes and frozen heart tissue sections. For immunofluorescence combined with OPP assay, immunostaining was performed first, followed by the OPP assay as described below. For WGA staining, sections underwent incubation with FITC conjugated WGA for 1 hour at RT (diluted 1:10 in PBSx1) prior to the permeabilization step.

### TUNEL Assay

The assessment of apoptosis was conducted using the Click-iT TUNEL Alexa Fluor 488 imaging assay (Thermo-Fisher Scientific) according to the manufacturer instructions. For the positive control, sections were treated with DNase I-containing solution for 30 minutes at RT prior to conducting the TUNEL assay.

### Rodents

Mice and rats were maintained in a pathogen free facility, temperature and humidity control, and a 12-hour light cycle with ad libitum water and food supply. Male and females were equally used.

### Study approval

All procedures and protocols were approved in advance by the Technion animal care and use committee (IACUC) and performed according to IACUC guidelines.

### Single-molecule FISH

Stellaris RNA FISH probes (Biosearch technologies) were used, tagged with either 5-TAMRA or Quasar-670 fluorophores. smFISH was performed according to the manufacturer’s protocol, as previously described (7). Slides were imaged with Axio Observer inverted fluorescent microscope (Zeiss, see under microscopy). Exposure times for smFISH signals were between 800-3000 milliseconds. Images were acquired as a full thickness z-stack with a 0.24 µm section thickness. Images were imported into the ImageJ software, a Laplacian of Gaussian filter was applied to the smFISH channels using the Log3D plugin, and a maximum intensity merge of the z-stack was performed.

### RT-qPCR

For cell cultures, RNA was isolated using the NucleoSpin RNA (Macherey-Nagel #740955) according to the manufacturer’s instructions. For isolation of RNA from heart tissues, atrial tissue was removed, and a Bullet Blender homogenizer (Next Advance) was used to homogenize the whole ventricular tissue prior to the standard protocol of RNA extraction. All studies were performed with a standard curve, technical duplicates or triplicates, and controls (negative control and no reverse transcriptase control).

### Polysome profiling

Whole hearts with atria removed were dissected into small pieces and homogenized with a Bullet Blender homogenizer (Next Advance) containing polysome extraction buffer (20 mM Tris-HCl PH 7.4, 200 mM KCl, 10 mM MgCl_2_, 1 mM DTT, 0.15 mg/mL Cycloheximide (Sigma Aldrich, 0180-5G), 1x Protease inhibitors cocktail). The homogenates were centrifuged at 17,000 g for 12 minutes at 4 °C. Triton X-100 was added to a final concentration of 1%, followed by 30 min incubation at 4 °C. The samples were centrifuged again and the supernatant was transferred to a pre-chilled new tube. Linear sucrose gradients of 10–50% were prepared in SW41 centrifugation tubes (Beckman Coulter, Ultra-Clear™ 344059), using the Gradient Station (BioComp). Gradient Solutions 1 (GS1) and 2 (GS2) were prepared with RNase/DNase-free UltraPure water and filtered with a 0.22 μm filter, consisting of 20 mM Tris–HCl pH 8, 200 mM KCl, 10 mM MgCl_2_, 0.2 mg/ml CHX and 10% or 50% w/v RNase-free sucrose, respectively.

SW41 centrifugation tubes were filled with GS1, followed by GS2 layered at the bottom of the tube. The linear gradient was formed using the tilted methodology, with the Gradient Station Maker (BioComp). 700 μL of each lysate was loaded on top of the gradients and centrifuged at 4 °C for 150 min at 35,000 rpm. Gradients were immediately fractionated using the Gradient Station, and 24 × 450 μL fractions were collected while monitoring the absorbance at 260 nm continuously. Proteins from individual fractions were extracted by adding freshly made Trichloroacetic acid (Carlo Erba, 411527) to a final concentration of 10% v/v, and allowed to precipitate overnight at −20 °C. Samples were then pelleted at 17,000 g for 15 min at 4°C, supernatant discarded, and the pellet washed in ethanol and air dried for 15 min. The pellets were re-suspended in Laemmli buffer, heated at 95°C for 5 min, and loaded on SDS-PAGE gels for subsequent western blot analysis.

### BioID Proximity Labeling Assay

NRVM were transduced using adenoviruses encoding either 3x-HA-Cypher1s-mycBioID2 (bait construct) or mycBioID2 (control construct) and incubated for 48 hours to allow for expression of the encoded constructs. Culture medium was supplemented with 50 µM Biotin and the cells were incubated for 18 hours to allow for labelling. Subsequently cells were scraped and pellet was then frozen in liquid nitrogen.

### Pull-down of biotinylated proteins

Pull-down procedure was based on a previously published protocol (32). The supernatant was incubated with Streptavidin magnetic beads (EMD Millipore). Beads were washed twice with wash buffer 1 (2% SDS), once with wash buffer 2 (0.1% Deoxycholic acid, 1% Triton X-100, 1 mM EDTA, 500 mM NaCl, 50 mM HEPES), once with wash buffer 3 (250 mM Lithium chloride, 10 mM Tris-HCl, 1 mM EDTA, 0.5% NP-40, 0.5% Deoxycholic acid) and three times with 50 mM Tris-HCl. Washed beads were resuspended in 50 mM Ammonium Bicarbonate.

### Mass spectrometry

The proteins on the beads were reduced in 3mM DTT, 100mM ammonium bicarbonate and 8M Urea, modified with 10mM iodoacetamide in 100mM ammonium bicarbonate and 8M Urea (room temperature 30 min in the dark) and digested in 2M Urea, 25mM ammonium bicarbonate with modified trypsin (Promega), overnight at 37°C in a 1:50 (M/M) enzyme-to-substrate ratio. The tryptic peptides were desalted using homemade C18 stage tip, dried and re-suspended in 0.1% Formic acid. The peptides were resolved by reverse-phase chromatography on 0.075 X 300-mm fused silica capillaries (J&W) packed with Reprosil reversed phase material (Dr Maisch GmbH, Germany). The peptides were eluted with linear 60 minutes gradient of 5 to 28% 15 minutes gradient of 28 to 95% and 15 minutes at 95% acetonitrile with 0.1% formic acid in water at flow rates of 0.15 μl/min. Mass spectrometry was performed by Q Exactive HF mass spectrometer (Thermo) in a positive mode (m/z 300–1800, resolution 60,000 for MS1 and 15,000 for MS2) using repetitively full MS scan followed by high collision induces dissociation (HCD, at 27 normalized collision energy) of the 18 most dominant ions (>1 charges) selected from the first MS scan. A dynamic exclusion list was enabled with exclusion duration of 20 s.

The mass spectrometry data was analyzed using the MaxQuant software 1.5.2.8 for peak picking and identification using the Andromeda search engine, searching against the rat proteome from the Uniprot database with mass tolerance of 6 ppm for the precursor masses and 20 ppm for the fragment ions. Oxidation on methionine and protein N-terminus acetylation were accepted as variable modifications and carbamidomethyl on cysteine was accepted as static modifications. Peptide- and protein-level false discovery rates (FDRs) were filtered to 1% using the target-decoy strategy. The mass spectrometry proteomics data were deposited to the ProteomeXchange Consortium via the PRIDE (33) partner repository with the dataset identifier PXD046385.

### Proteomic analysis

Proteomic analysis was performed using MaxQuant and Perseus with log2 of LFQ values. Proteins that were identified by less than two distinct peptides were removed. In addition, a protein was considered for the dataset and for further analysis if it had a valid value in two or more replicates in at least one of the groups (control or experiment). Imputation of missing values was performed using Perseus by random numbers drawn from a normal distribution. We used a cutoff of log2FC≥1 and t test p<0.05 for significantly enriched proteins. The R EnhancedVolcano package (Blighe, K, S Rana, and M Lewis. 2018. “EnhancedVolcano: Publication-ready volcano plots with enhanced colouring and labeling.”) was used to draw volcano plots. The ShinyGO 0.77 tool was used for pathway enrichment (34). STRING was used for visualization (35).

### OPP assay

Analysis of the location of protein synthesis was performed using with o-propargyl-puromycin (OPP) and the Click-iT Plus OPP Alexa Fluor 488 Protein Synthesis Assay Kit (Thermo-Fischer Scientific) as previously described (7). For in-vivo protein synthesis assay mice were injected intraperitoneally with OPP (0.049 mg/gr). Mice were sacrificed one hour following OPP injection. The heart was then removed, frozen and cryosectioned.

### Statistics

Statistical methods are described in each of the figure legends. P values <0.05 were considered significant.

## Data availability

The mass spectrometry proteomics data were deposited to the ProteomeXchange with the dataset identifier PXD046385. Values for all data points in graphs are reported in the Supporting Data Values file.

## Author contributions

**IK** conceived of and conceptualized the study, designed experiments, wrote the manuscript, and interpreted the data with intellectual input from all authors. **RH**, **OS**, **TZ**, and **LC** conceptualized and designed experiments, conducted and analyzed experiments, and contributed to the review and editing of the manuscript. **IE** developed tools and performed analysis. **JV** contributed resources and contributed to the review and editing of the manuscript. **AS** Developed tools for the analysis of the data and contributed to the review and editing of the manuscript.

## Acknowledgements

Funding for this work was provided by the Israel Science Foundation (grant # 1385/20) and by the Foundation Leducq Research grant no. 20CVD01 to IK and JvdV The authors want to thank the Biomedical Core Facility at the Faculty of Medicine, Technion; the Pre – Clinical Research Authority at the Technion, and the Smoler Proteomics Center, Technion-Israel Institute of Technology

---

**Supplemental Figure 1.**
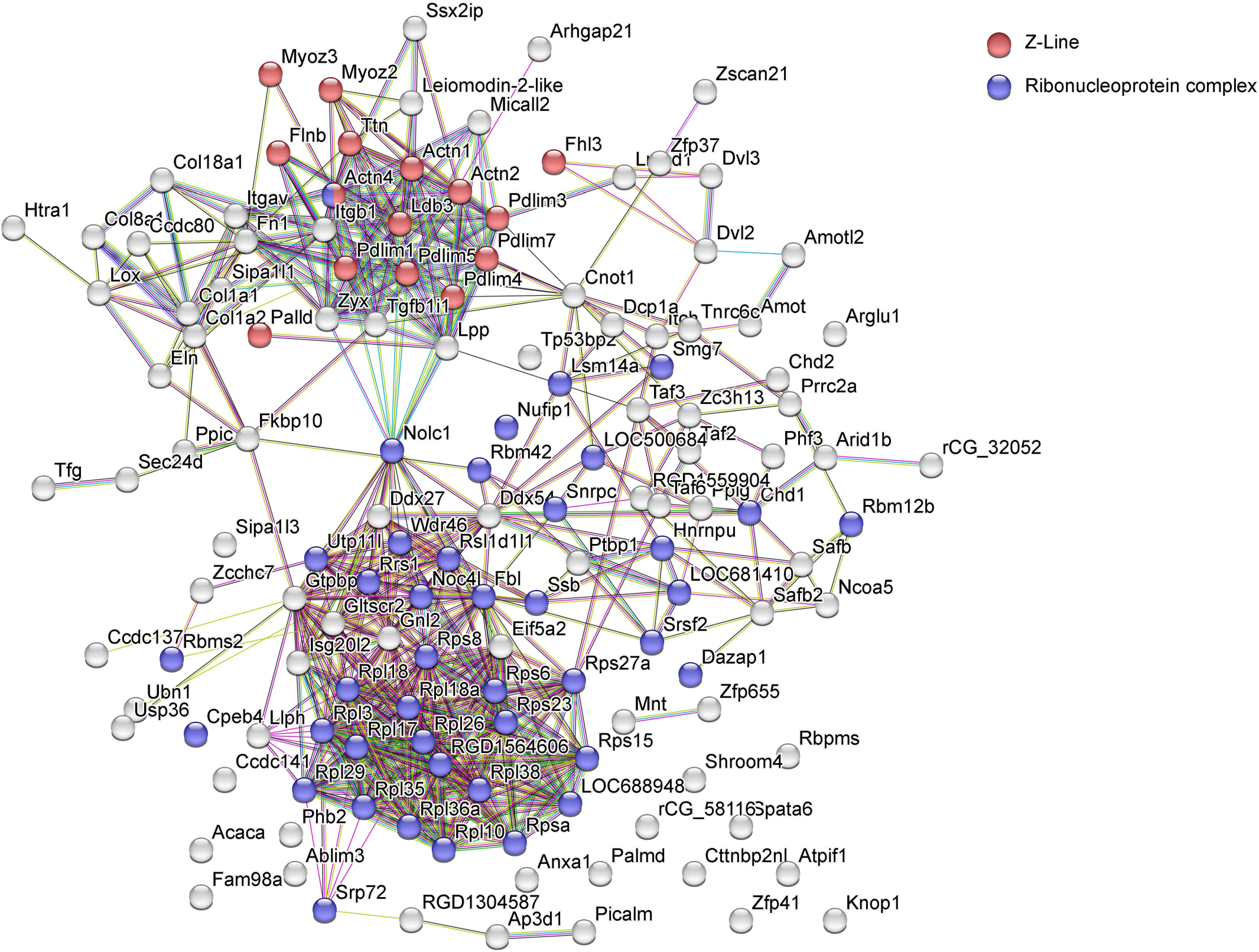
STRING of Cypher-BioID2 identified proteins shows Z-line proteins and components of the translational machinery. STRING protein-protein interaction map of proteins significantly enriched by Cypher-BioID2. Proteins labeled in red belong to the Z-line GO term (GO:0030018), FDR for enrichment = 2.61e-11. Proteins labeled in purple belong to the Ribonucleoprotein complex GO term (GO:1990904), FDR for enrichment = 3.83e-16.

**Supplemental Figure 2.**
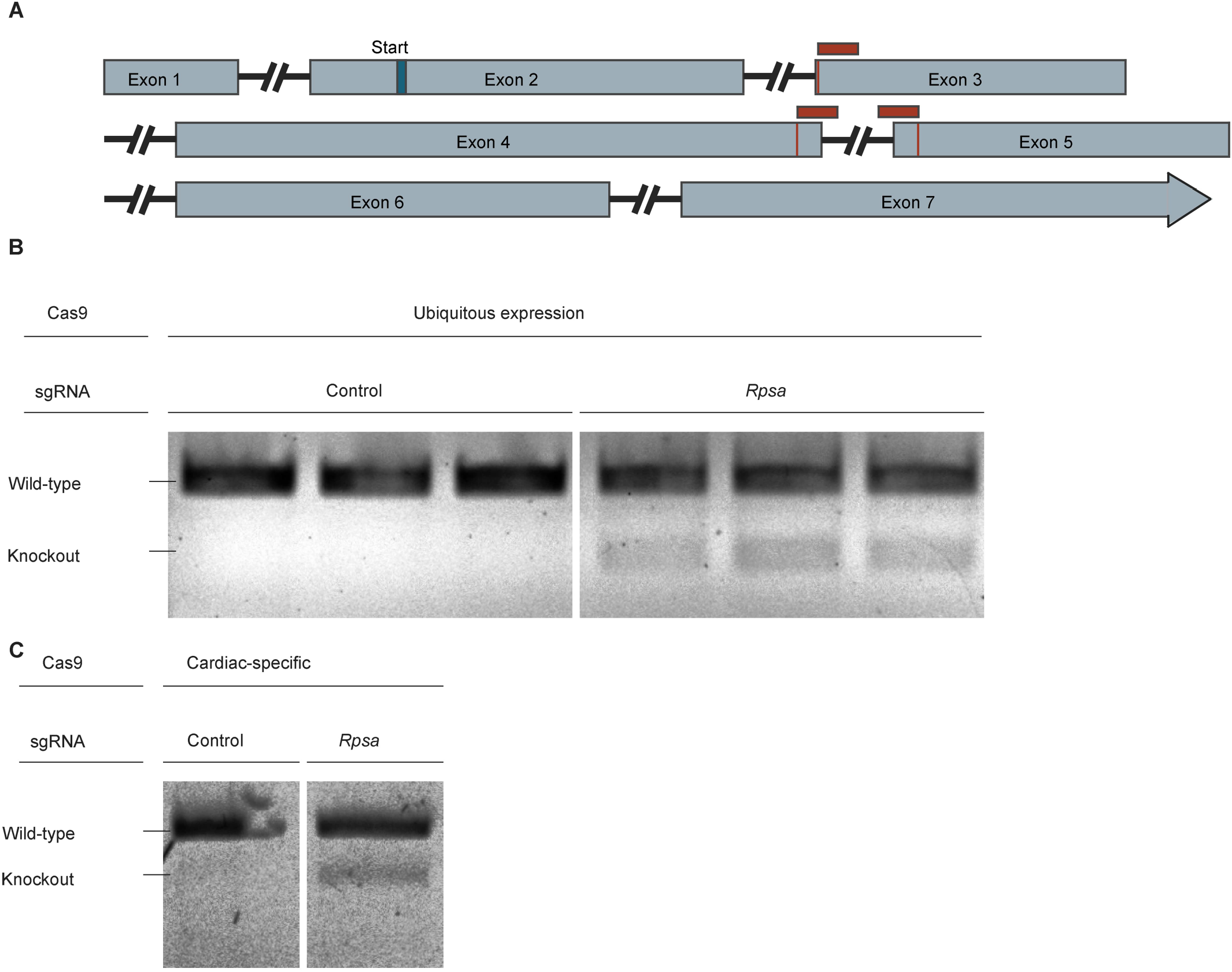
Validation of CRISPR-CAS9 knockout of *Rpsa* in vivo. (**A**) Illustration of the *Rpsa* gene with location of sgRNAs targeting exons 2, 3 and 4 (red) and predicted cut sites. (**B** and **C**) T7 Endonuclease 1 (T7E1) assay in mouse heart genomic DNA PCR amplicon surrounding the expected CAS9 cut site in *Rpsa* gene showing successful editing with indels in the *Rpsa* gene in knockout mice. In these mosaic mice we find both edited and unedited cells. The upper band indicates the presence of the wild-type *Rpsa* and the bottom band indicates *Rpsa* with indels. This lower band is observed only in knockout mice transduced with *Rpsa* sgRNA but not with control sgRNA. (**B**) Genomic DNA from control and knockout mice with unconditional CAS9 expression, and (**C**) with DNA from Lox-stop-Lox-Cas9 mice transduced with troponin driven CRE AAV.

**Supplemental Figure 3.**
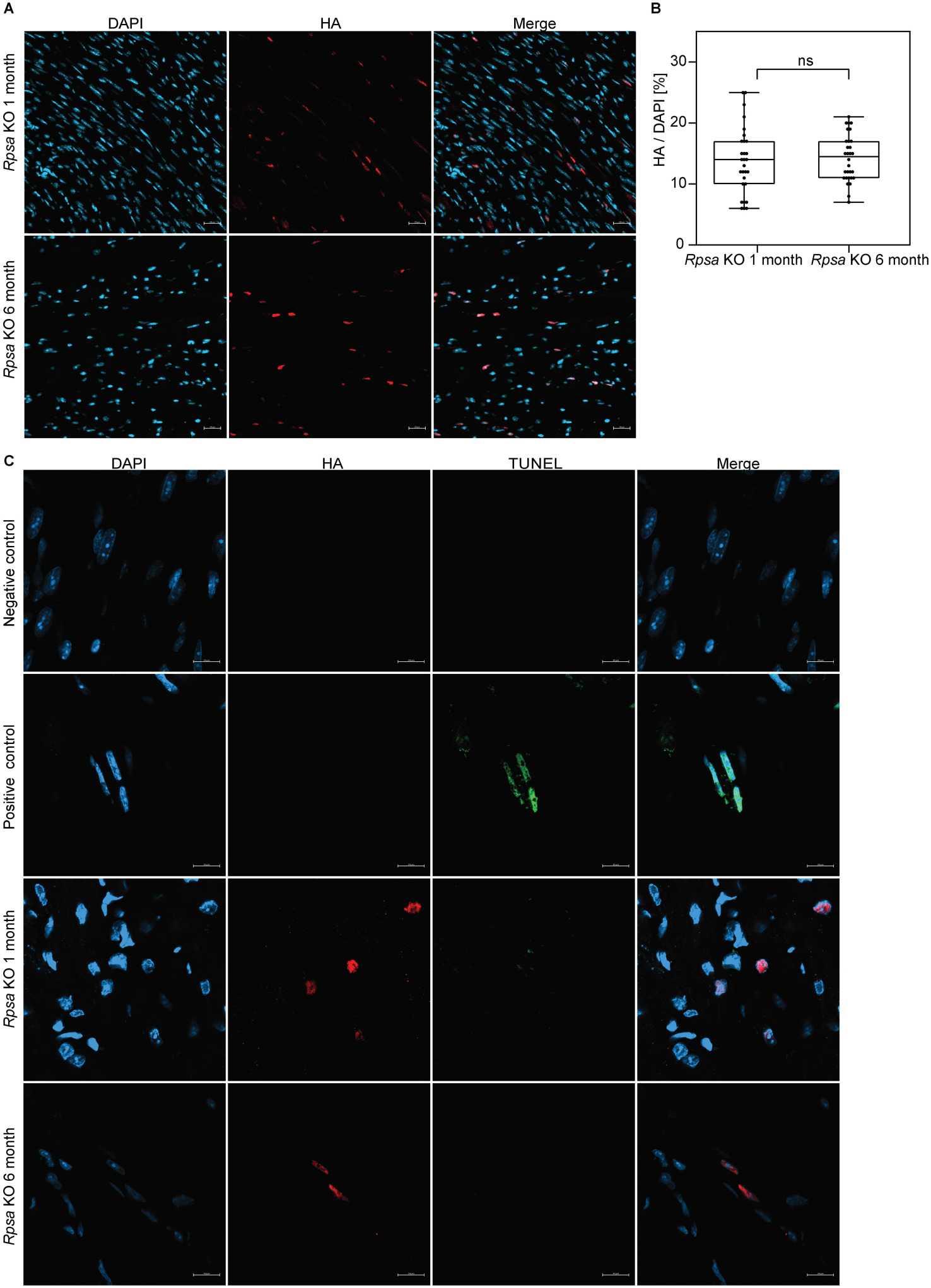
Analysis of targeted cell numbers and apoptosis in *Rpsa* knockout hearts. (**A** and **B**) Representative images and quantitative analysis of targeted HA-CRE (red) percentage in *Rpsa* knockout hearts, at 1 and 6 months of age showing no statistically significant difference, nuclei were counter stained with DAPI (blue). Scale bar = 20μm. N = 2-3 mice per experimental group, n = 29-30 analyzed images per group. ns (p > 0.05), by Student’s t test. Data are presented as individual values with box plot displaying the median with 25th and 75^th^ percentiles. (**C**) Representative images of TUNEL staining of apoptosis. No positive TUNEL (Green) nuclei were detected in *Rpsa* knockout mice, at either 1 or 6 months of age. Positive control is DNase I treated section. Scale bar = 10μm. N = 2-3 mice per group.

**Supplemental Figure 4.**
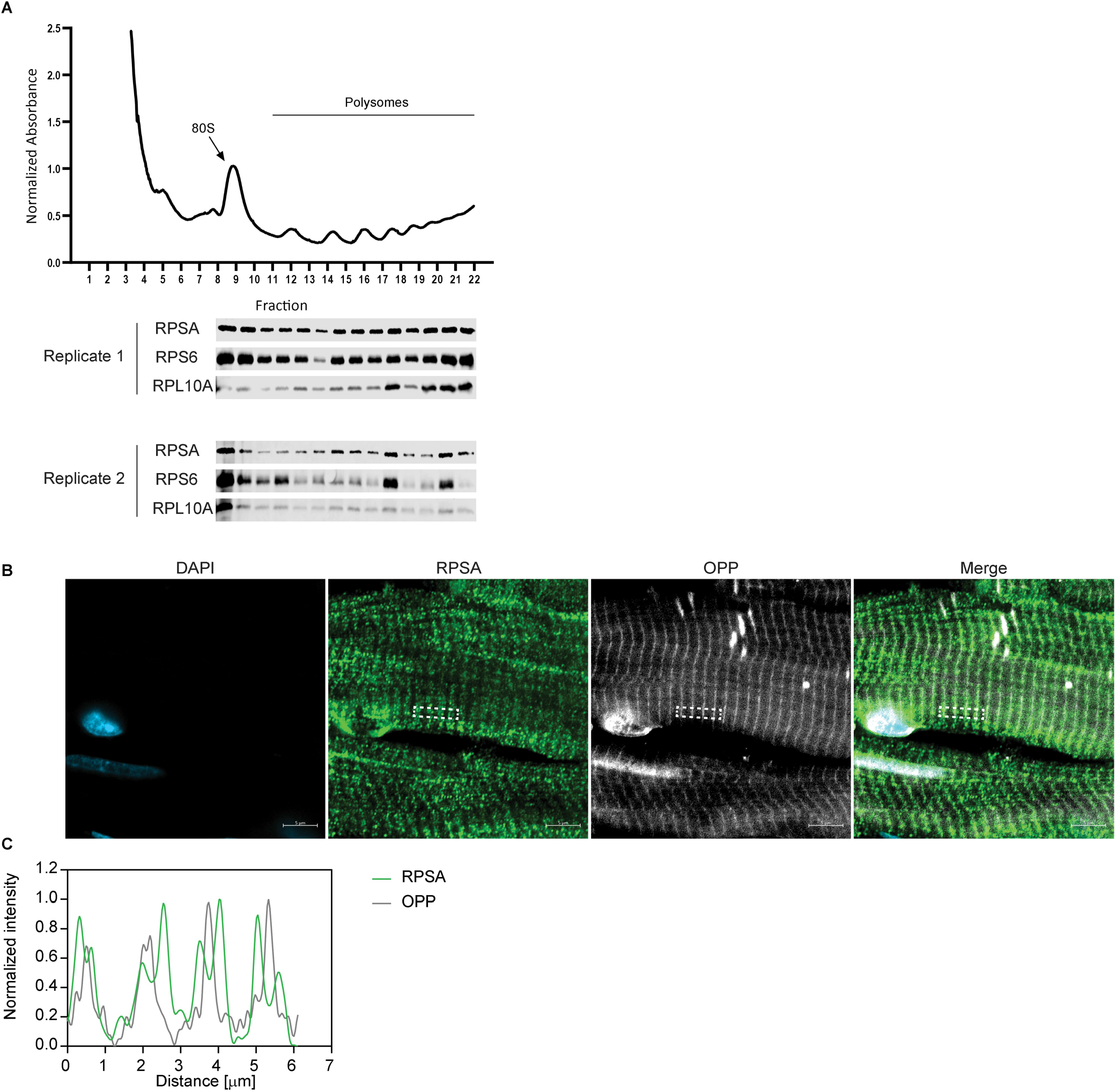
RPSA is associated with translating ribosomes. (**A**) Polysome profile and Western blot analyses of normal mosue heart. Western blot of monosome (80S) and polysome fractions were probed with anti-RPSA, anti-RPS6, and anti-RPL10A antibodies showing association of RPSA with translating ribosomes. The experiment was performed in two biological replicates. (**B**) Immunofluorescence images of cardiac sections from mice injected with OPP and stained for OPP (white) and RPSA (green). OPP labeled nascent proteins have a sarcomeric Z-line striated pattern, and RPSA resides in close proximity on both sides of the OPP signal. Scale bar = 5 μm. (**C**) Line scan analysis depicting the prominent, close proximity of RPSA at both sides of OPP labeled nascent proteins.

**Supplemental Figure 5.**
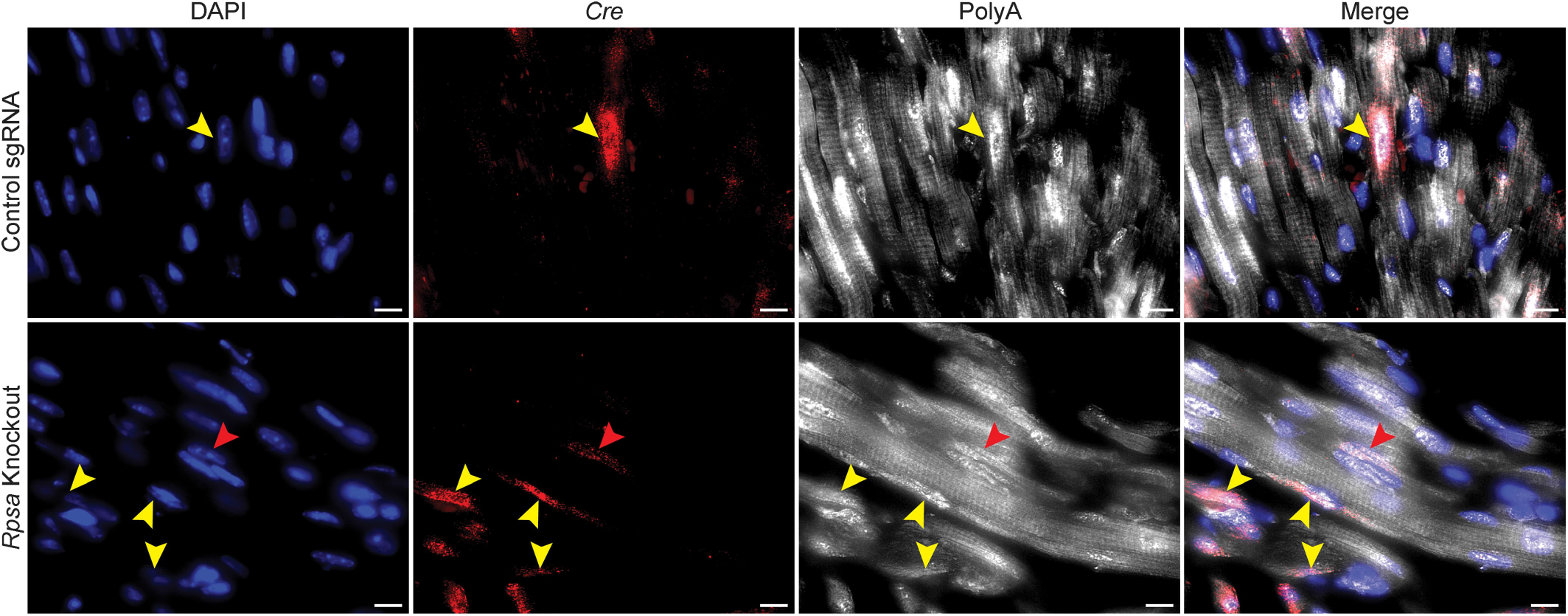
***Rpsa* Knockout effect on mRNA distribution.** Representative dual smFISH images from *Rpsa* knockout and control hearts incubated with *Cre* (red) and *polyA* (white) probes. In untargeted cells and control sgRNA hearts, the *polyA* mRNA signal exhibits a cross-striated pattern. *Rpsa* targeted cardiomyocytes appear smaller yet show cytoplasmic *polyA* signal. Despite their reduced size, *polyA* striations can be observed in some cells (red arrowhead). Yellow arrowheads indicate *Cre*-positive cells.

**Supplemental Figure 6.**
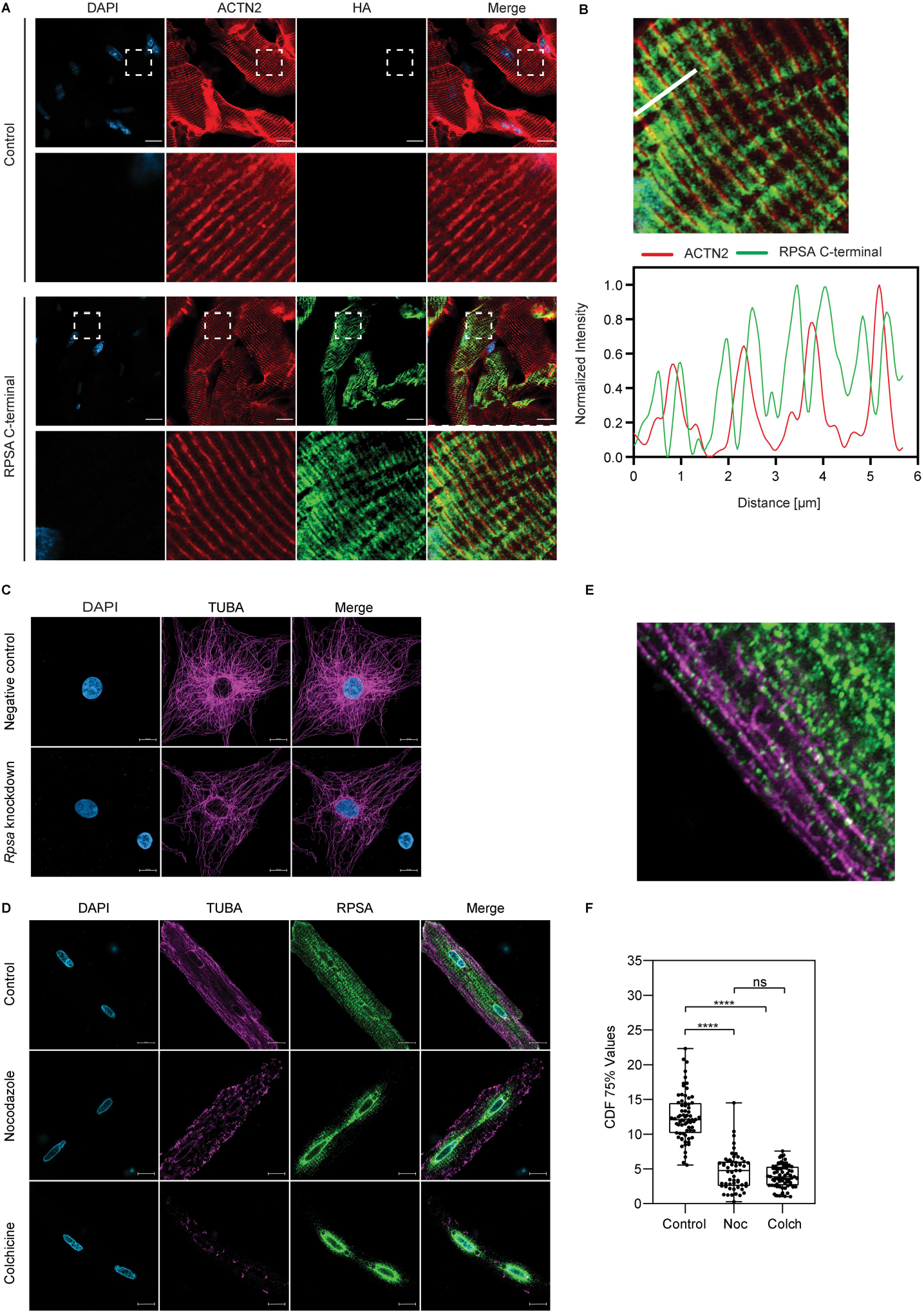
RPSA Z-line localization depends on C-terminal fragment and microtubules. (**A** and **B**) Representative images and line scan analysis of cardiac sections from mice transduced with AAV encoding for HA-tagged mCherry-RPSA C-terminal domain. Staining with antibodies for HA (pseudo-colored green) and α-Actinin (pseudo-colored red) shows that the C-terminal domain of RPSA is sufficient for localization to both sides of the Z-line, where endogenous RPSA and where ribosomes are localized. (**C**) Immunofluorescence images for NRVMs stained for Tubulin (TUBA), purple, following *Rpsa* siRNA mediated knockdown or control siRNA (Negative control). While *Rpsa* knockdown did not alter the reticular microtubule pattern, a reduction in total TUBA signal was observed. N = 2 biological replicates. (**D**) Representative images of adult rat cardiomyocytes following microtubule ablation by Nocodazole (Noc) or Colchicine (Colch) showing loss of RPSA’s cross-striated localization pattern and a shift to a peri-nuclear localization. (**E**) Higher magnification of control cell from D showing intersection of microtubules and RPSA. (**F**) For quantification of RPSA distribution we used the CDF75% measuring the distance in μm from the nucleus in which 75% of the RPSA signal was found, showing peri-nuclear collapse with microtubule ablation. N= 3 biological replicates. N = 55-71 analyzed cell per group. ns (p > 0.05), **** (p ≤ 0.0001), by 1-way ANOVA test. Scale bar = 10μm. Data are presented as individual values with box plot displaying the median with 25th and 75^th^ percentiles.

**Supplemental Figure 7.**
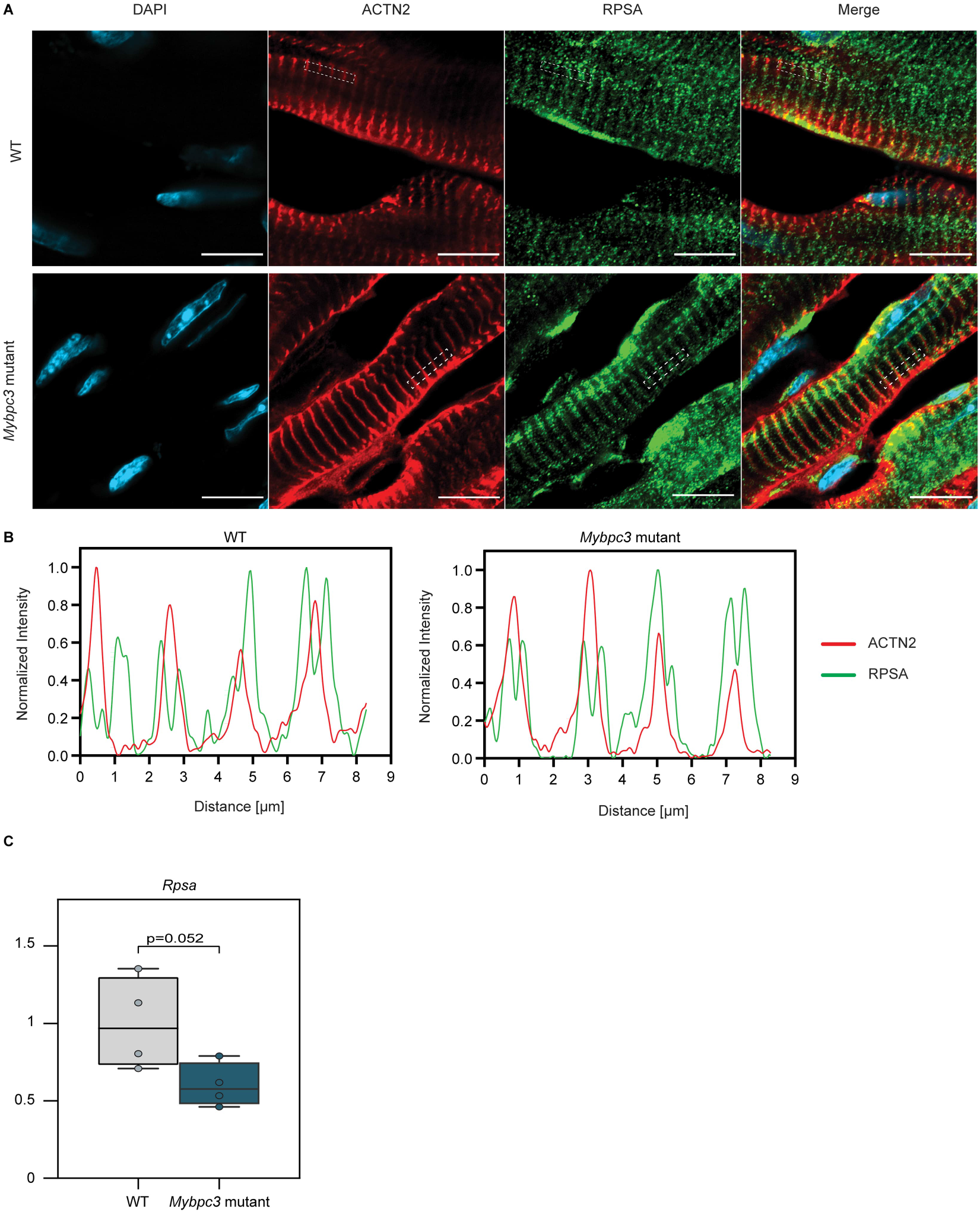
The subcellular localization of RPSA and its mRNA expression in *Mybpc3* mutant mice. (**A** and **B**) Representative immunofluorescence images and line scan analysis in Wild Type (WT) and *Mybpc3* mutant mice show unaltered RPSA localization pattern on both sides of the Z disc. RPSA (green), ACTN2 (red), nuclei counterstained with DAPI (blue). N = 3 mice per group. Scale bar = 10μm. (**C**) RT-qPCR analysis of *Rpsa* gene expression in WT and *Mybpc3* mutant mice shows a trend towards reduced *Rpsa* expression (Student’s t test, p = 0.052). Expression data was normalized to *Gapdh*. n=4 mice per group. Data are presented as individual values with box plot displaying the median with 25th and 75^th^ percentiles.

**Supplemental video 1.** Three dimensional video representation of isolated adult rat ventricular cardiomyocyte stained for α-actinin (ACTN2) (red), RPSA (green) and DAPI (blue) for nuclei, illustrating the sarcomeric localization of RPSA in cardiomyocyte. Z-stack images were acquired using Ziess LSM900 inverted confocal microscope, and image processing with 3D reconstruction was performed using Imaris software.

